# Manipulation of RNA Polymerase III by Herpes Simplex Virus-1

**DOI:** 10.1101/2021.07.21.453214

**Authors:** Sarah E. Dremel, Frances L. Sivrich, Jessica Tucker, Britt A. Glaunsinger, Neal A. DeLuca

## Abstract

RNA Polymerase III (Pol III) transcribes noncoding RNA, including transfer RNA (tRNA), and acts as a pathogen sensor during the innate immune response. To promote enhanced proliferation, the Pol III machinery is commonly targeted during cancer and viral infection. Herein we employ DM-RNA-Seq, 4SU-Seq, ChIP-Seq, and ATAC-Seq to characterize how Herpes Simplex Virus-1 (HSV-1) perturbs the Pol III landscape. We find that HSV-1 stimulates tRNA expression 10-fold, with mature tRNAs exhibiting a 2-fold increase within 12 hours of infection. Perturbation of host tRNA synthesis requires nuclear viral entry, but not synthesis of specific viral transcripts, nascent viral genomes, or viral progeny. Host tRNA with a specific codon bias were not targeted—rather increased transcription was observed from euchromatic, actively transcribed loci. tRNA upregulation is linked to unique crosstalk between the Pol II and III transcriptional machinery. While viral infection is known to mediate host transcriptional shut off and lead to a depletion of Pol II on host mRNA promoters, we find that Pol II binding to tRNA loci increases. Finally, we report Pol III and associated factors bind the HSV genome, which suggests a previously unrecognized role in HSV-1 gene expression. These data provide insight into novel mechanisms by which HSV-1 alters the host nuclear environment, shifting key processes in favor of the pathogen.

## INTRODUCTION

Herpes Simplex Virus-1 (HSV-1) is a ubiquitous human pathogen which most commonly causes recurrent lesions of the oral and genital mucosa. The virus is associated with a wide range of additional pathologies—including herpes keratitis, herpetic whitlow, and encephalitis, to name a few—representative of the large range of cells permissive to replication. Similar to other herpesviruses, HSV-1 establishes a latent reservoir in the peripheral nervous system and reactivates to cause disease in response to various physiological stimuli.

HSV-1 replicates and assembles almost entirely within the host nucleus, reprogramming the host transcriptional machinery to prioritize expression of ∼90 viral messenger RNAs (mRNAs) (1). These genes are transcribed in a temporally coordinated sequence, such that their protein products are expressed at the appropriate time in the life cycle of the virus (2, 3). Immediate early (IE or *α*) gene products enable the efficient expression of early (E or *β*) and late (L or *γ*) genes. The protein products of E genes are mostly involved in DNA replication. DNA replication and IE proteins enable the efficient transcription of L genes, which encode the structural components of the virus. DNA replication licenses L promoters, enabling the binding of core Pol II transcription factors, thus activating the initiation of L transcription (4). Productive HSV-1 infection is incredibly rapid, causing a single infected cell to produce progeny between 4 and 6 hours (h) postinfection, culminating in ∼1,000 infectious progeny within 18 h. Considering the rapid HSV-1 life cycle, it is perhaps unsurprising that within 6 h of infection, viral transcripts rise to 50% of the total mRNA within a host cell. The corresponding decrease in host transcripts is facilitated by two mechanisms i. VHS-mediated mRNA decay (5-7) and ii. ICP4-DNA mediated decrease of Pol II on mRNA promoters (8-11).

Transcriptional studies of HSV center on mRNA and RNA Polymerase II (Pol II), and thus ignore the potential contributions of other DNA-dependent RNA Polymerases, namely Pol I and Pol III (12, 13). Pol I transcribes a single multi-copy transcript, 45S, which is spliced and processed to produce 5.8S, 18S, and 28S ribosomal RNA (rRNA). Pol III transcribes noncoding RNAs, including 5S rRNA, transfer RNA (tRNA), Alu elements, 7SL, 7SK, U6, and select microRNA (miRNA). Pol III also transcribes noncoding RNAs (ncRNA) for various DNA viruses, including: VAI and VAII RNAs of adenovirus (14), EBER1 and EBER2 of Epstein Barr Virus (15) and the tRNA-miRNA encoding RNAs (TMERs) of MHV68 (16-18).

In this study, we explore HSV-1 alteration of the host Pol III transcription landscape. Pol III transcription requires its own unique set of general transcription factors, and the basal promoter requirements have been classified into three different promoter types (19). Type I promoters include 5S rRNA genes, and use the TFIIIB complex (BDP1, TBP, BRF1), TFIIIC complex (GTF3C1, GTF3C2, GTF3C3, GTF3C4, GTF3C5, GTF3C6), and TFIIIA (GTF3A). Type II promoters include tRNAs and require the TFIIIB and TFIIIC complexes. Type III promoters include tRNA-Sec, U6 and 7SK and use distal enhancers including STAF, OCT1, SNAP as well as a distinct TFIIIB complex (BDP1, TBP, BRF2). Each of these promoters consists of distinct combinations of internal cis-acting sites (A, B, C, IE). To add another layer of complexity, recent studies have demonstrated crosstalk between Pol II and Pol III. Highly expressed Pol III transcripts are located in regions of open chromatin adjacent to Pol II promoters (20). Additionally, Pol II binding is observed at highly expressed Pol III promoters (21), and Pol III occupancy frequently scales with nearby levels of Pol II (19, 22).

Viruses have evolved unique mechanisms to invade hosts, alter cellular pathways, and redirect host factors for viral processes. A number of DNA viruses, including adenovirus (23-25), SV40 (26), and Epstein Barr Virus (27), have been shown to increase Pol III transcription in infected cells. These viruses employ a viral regulatory protein (E1A, E1B, T-antigen, and EBNA1) to mediate increase of Pol III transcription factor abundance. Additionally, infection with the gammaherpesvirus MHV68 has been linked to increased expression of murine B2 SINE retrotransposons and pre-tRNAs, suggesting that viral manipulation of Pol III may impact both viral and host Pol III transcription (28, 29). HSV-1 has previously been shown to induce the Pol III type II transcript, Alu repeat units (30, 31). How HSV-1 affects other Pol III-dependent transcripts and the mechanism behind Alu upregulation is yet unknown.

Herein we report a comprehensive characterization of how HSV-1 alters the Pol III transcriptional landscape, ultimately increasing the pool of tRNA available during productive infection. Changes in tRNA levels coincided with increased Pol II recruitment at these loci. This discovery is at odds with the general environment of Pol II depletion from host mRNA promoters which contributes to host transcriptional shut off (8-11). We also report, for the first time, recruitment of Pol III to the HSV-1 genome. Whether this binding event results in production of a Pol III-dependent transcript is still unknown. However, these results suggest a novel role of Pol III in promoting and regulating HSV-1 transcription.

## RESULTS

### Impact of HSV-1 Productive Infection on Pol I and III Transcripts

We began by assessing changes in noncoding RNA species, particularly those expressed by Pol I and III. Human diploid fibroblast (MRC5) cells were mock-infected, infected with ΔICP0/4/22/27/47 (d109), ΔICP4 (n12), or wildtype (KOS) HSV-1. d109 infection lacks synthesis of all viral proteins and viral genomes; however, it robustly stimulates a cGAS-mediated innate immune response (32, 33). n12 infection is deficient in the synthesis of early (E) and late (L) viral proteins, nascent viral genomes, and viral progeny (34). We observed little change in Pol I transcripts 18S rRNA, 45S pre-rRNA, or 5.8S rRNA using RT-qPCR (Fig. 1A) and northern blot analysis (Fig. 1B). Pol III type I, II, and III transcripts were affected disparately (Fig. 1A-B). Type I transcript 5S rRNA was unaltered by infection, while type II transcripts were strongly upregulated. Infection with n12 and wildtype HSV-1 increased the levels of type II transcripts from 2 to 16-fold (Fig. 1A-C). This upregulation was most drastic for pre-tRNA, and specific to tRNA with type II promoters as we did not observe upregulation for a tRNA encoded by a type III promoter, namely tRNA selenocysteine (tRNA Sec). We observed an increase—albeit modest ∼1.5-fold—in mature tRNA (Fig. 1C). Type III transcripts U6 snRNA, tRNA-sec, and 7SK decreased minimally during infection (Fig. 1A-B). n12 infection phenocopied transcriptional changes in wild-type infection, indicating that synthesis of E and L viral proteins, nascent viral genomes, or viral progeny were not required for the virus to alter the Pol III transcriptional landscape. In contrast, d109 infection had no impact on the transcriptional landscape, indicating viral nuclear entry or the induction of an innate immune response is not sufficient to induce tRNA upregulation. These results demonstrate HSV-1 selectively targets and upregulates Pol III type II transcripts.

**Fig. 1.**
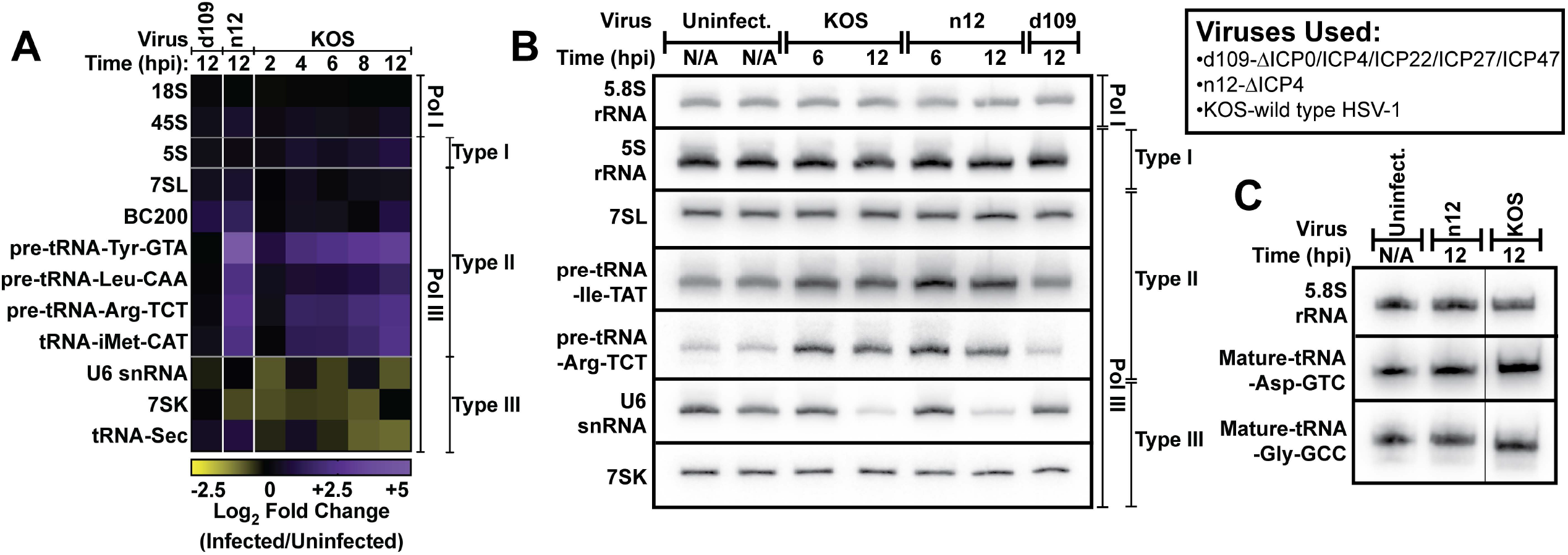
Impact of HSV-1 Productive Infection on Pol I and III Transcripts. Human fibroblast cells were mock-infected or infected with d109, n12, or wild-type HSV-1 (KOS). RNA was isolated at indicated times, and A) RT-qPCR or B-C) northern blots were used to assess transcript abundance. A) Data is the average of biological triplicate experiments, and values are average log_2_ fold change of infected over uninfected samples. cDNA copy number was calculated as a function of total RNA used for reverse transcription. B-C) Images are representative northern blots from biological duplicate experiments.

### Alteration of the tRNA landscape by HSV-1 Productive Infection

To investigate global changes in tRNA species during HSV-1 productive infection, we performed DM-tRNA-Seq (35). Cells were mock-infected or infected with n12 or KOS for 12 hours. Using this technique, we can accurately discriminate between pre-and mature-tRNA species and between the approximately 500 different tRNA-encoding loci within the host genome. We further delineated tRNA expression by those encoded from the nuclear or mitochondrial (MT) host genome.

We found that the total amount of nuclear-encoded tRNA increased 2-fold after infection with wildtype or ΔICP4 HSV-1 (n12). Pre-tRNA species were more affected than mature-tRNA, increasing 4 and 1.5-fold, respectively (Fig 2A). Nuclear-encoded tRNA expression was altered similarly between KOS and n12 infection (Fig. 2A-C), indicating only early viral life cycle events were required for the phenotype. In contrast, MT-encoded tRNA decreased ∼4-fold only in wildtype HSV-1 infection (Fig. 2A). We expected this decrease as HSV-1 degrades the host mitochondrial genome during productive infection (36). MT-tRNA changed minimally in n12 infection likely due to the absence of UL12 expression, which is the viral endonuclease responsible for MT-genome degradation.

**Fig. 2.**
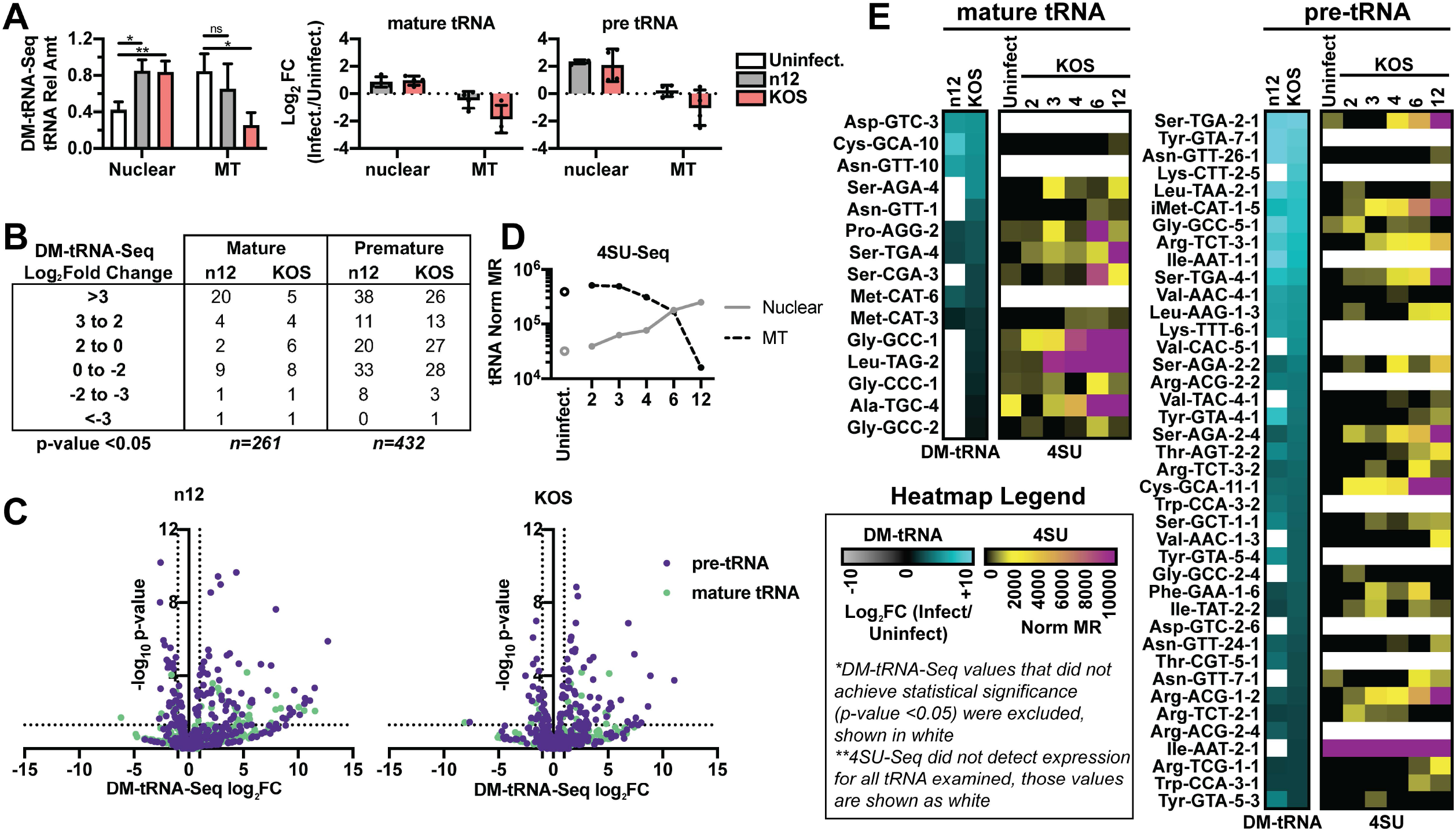
Alteration of the tRNA landscape by HSV-1 Productive Infection. Human fibroblasts were mock-infected or infected with ΔICP4 (n12) or wildtype HSV-1 (KOS) and RNA was isolated for DM-tRNA-Seq or 4SU-Seq. We used the tRNA assembly for hg38 predicted by GtRNAdb for mapping pre-or mature nuclear and mitochondrial (MT)-encoded tRNA. A, B, C, E) For DM-tRNA-Seq, RNA was collected 12 hpi and data was normalized to an internal spike-in control and the size in kb of each tRNA. A) Statistical values were generated from a paired t-test, where * indicates a p-value of <0.05, or ** <0.01. B-C) Differential expression analysis values are log_2_ fold change (infected/uninfected) versus the false-discovery rate (FDR) p-value. D-E) 4SU was pulsed in for 15 minutes at the time (hpi) indicated to label nascent RNA. Data was normalized to rRNA mapped reads and the size in kb of each tRNA. E) Heatmaps for upregulated mature-and pre-tRNA. Differentially expressed (DE) mature-tRNA were defined as p-value <0.05, log_2_ fold change (KOS/uninfected) >0.5. DE pre-tRNA were defined as p-value <0.05, log_2_ fold change (KOS/uninfected) >2. Data with a p-value >0.05 was not plotted and is white in the heatmaps.

Since pre-tRNAs were more upregulated than the mature form we hypothesized that infection increased nascent transcription of nuclear-encoded tRNA. To assess this, we pulsed in 4-thiouridine (4SU) for 15 minutes at various stages of infection before isolating RNA and performing 4SU-Seq. tRNA are very stable ncRNAs, with half-lives of about 3 days. Using 4SU-Seq we were able to assess how nascent transcription shifts, without being confounded by tRNA made prior to infection. Echoing our DM-tRNA-Seq results, nuclear-encoded tRNA increased and MT-encoded tRNA decreased after infection (Fig. 2D). Nascent nuclear tRNA levels were increased ∼10-fold at 12 hpi (Fig. 2D). In Fig. 2E we show data for the top differentially expressed (DE) mature (p-value <0.05, log_2_ fold change KOS/uninfected >0.5) and pre-(p-value <0.05, log_2_ fold change KOS/uninfected >2) tRNA species in the DM-tRNA-Seq dataset (Supplemental Table 1). We observed an increase in nascent tRNA detected for these DE mature-and pre-tRNAs (Fig. 2E).

We next assessed whether there was enrichment for select isodecoders among the tRNA species altered by infection (Fig. S2-1). The HSV-1 genome is unusually GC-rich, ∼68%, which means the viral coding sequence relies more heavily on tRNA with GC-rich anticodons (Fig. S2-1B). Our breakdown of DE tRNA found that the isodecoders altered did not target only GC-rich species (Fig. S2-1). Generally, altered mature and pre-tRNA loci isodecoders appeared random (Fig. S2-1C). Additionally, we did not observe a shift in the identity of tRNA expressed, as silenced tRNA loci were not suddenly expressed or vice versa (Fig. S2-1C). Further study of the actual codon usage truly sampled during infection would need to be determined, this was merely a theoretical codon usage assuming each HSV-1 protein was synthesized in the same amount. Additional work is necessary to determine which, if any, of the differentially expressed tRNAs are rate limiting in viral translation.

Thus far we have focused on upregulated tRNA, however there was a smaller subset of tRNA (n=32 pre-tRNA, n=10 mature tRNA) downregulated by HSV-1 infection (Fig. 2B-C). We evaluated the genomic position of up-and down-regulated tRNA and found that downregulated tRNA loci were located in close proximity to Pol II gene promoters (Fig S2-2). ∼50% of downregulated tRNA loci were located within 3 kbp of a protein-coding gene promoter, whereas only ∼15% of upregulated tRNA loci or all tRNA loci were promoter-adjacent (Fig S2-2D). These data suggest downregulated tRNA targets may be linked to the absence of host transcription at adjacent mRNA promoters. Based on these results it is unlikely that tRNA upregulation is due to Pol II-transcriptional run off or -transcriptional interference.

### Defining HSV-1 processes required for tRNA upregulation

Our results in Fig. 1 limit the viral processes which may be critical for tRNA upregulation, including: i. immediate early (IE) proteins other than ICP4, ii. viral ncRNA, or iii. amount, but not identity, of viral transcripts. To discriminate between these hypotheses, we infected MRC5 cells with various HSV-1 mutants and assessed tRNA abundance by Northern Blot (Fig. 3). Mutants for the viral IE proteins—n12 (ICP4), 5dl1.2 (ICP27), n199 (ICP22), n212 (ICP0)—induced pre-tRNA-Ile indicating that these proteins are not themselves responsible for tRNA upregulation (Fig. 3A-B). Surprisingly, infection with hp66, a mutant defective for genome replication, did not induce host tRNA (Fig. 3A-B). This result was phenocopied in cells treated with specific inhibitors of HSV-1 genome replication (phosphonoacetic acid and acyclovir) and infected with wildtype HSV-1 (Supplemental Fig. 3-1C). Since other mutants in the panel were also defective for viral genome replication, this life stage could not be responsible for induction of host tRNA (Fig. 3C). Results for other mutants in the panel determined that synthesis of viral L proteins, genome replication and virion assembly were not processes required for induction (Fig. 1, 3A-C).

**Fig. 3.**
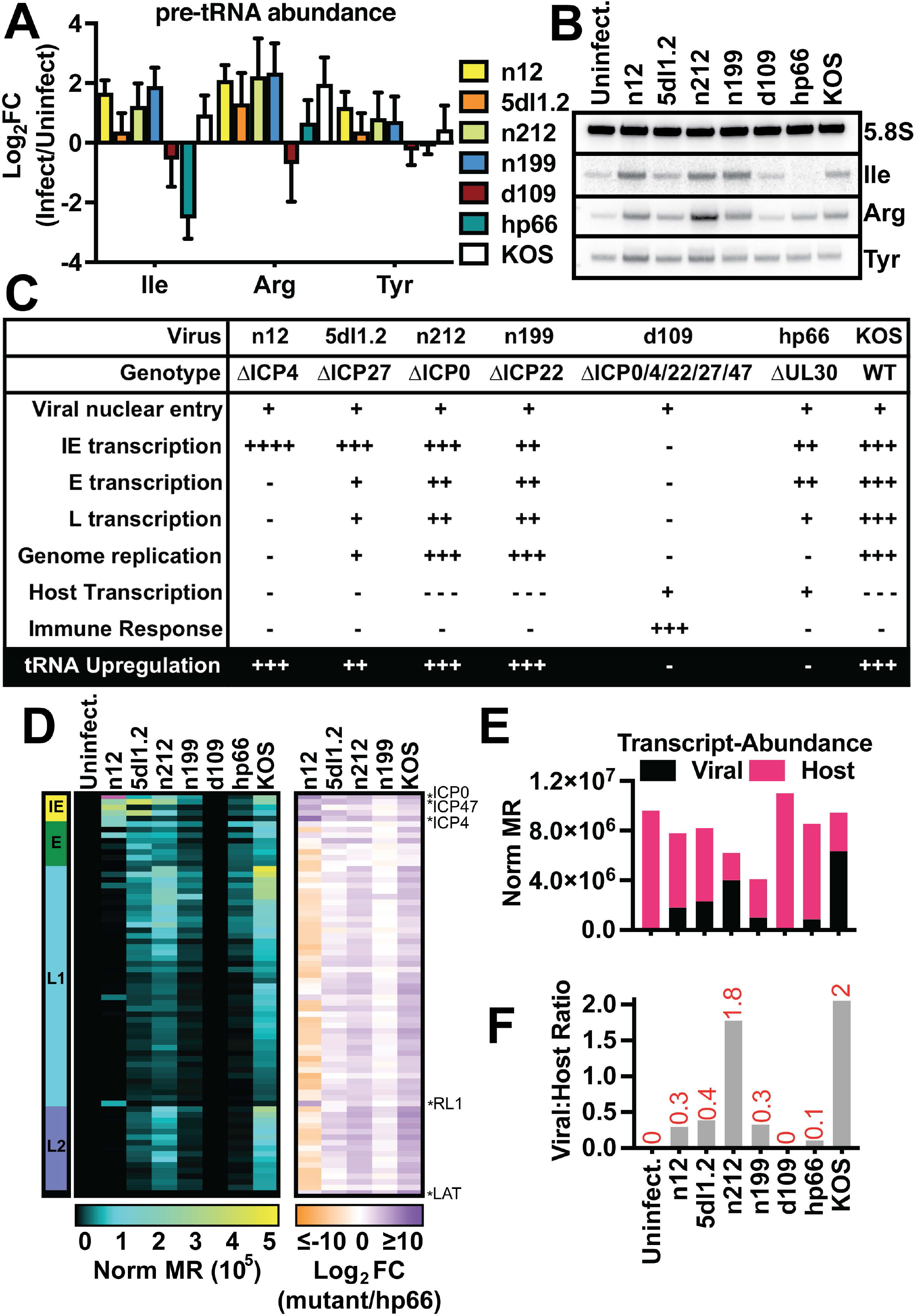
Defining HSV-1 processes required for tRNA upregulation. Human fibroblast cells were mock-infected or infected with n12, 5dl1.2, n212, n199, d109, hp66, or wild-type HSV-1 (KOS). RNA was isolated at 12 hpi, and A-B) Northern blots, or D-F) Ribo-depleted Total RNA-Seq was used to assess transcript abundance. A) Data is the average of biological replicates, error bars are standard deviation. B) Representative northern blot image of the data quantified in A. C) Summary of phenotypic differences between HSV-1 mutants used. D) Heatmaps for viral transcripts clustered by kinetic class with yellow as immediate early, green as early, blue as leaky late (L1), purple as true late (L2), and black as latency (LAT). Differentially expressed transcripts are highlighted with asterisks. E, F) All RNA-Seq reads mapping to the host or viral genome assembly plotted as abundance (million ERCC spike-in reads per kbp, Norm MR) or as the ratio of normalized viral transcripts: normalized host transcripts (Viral:Host Ratio).

We performed Ribominus-RNA-Seq on these samples to assess differentially expressed viral genes (Fig. 3D-F). We employed ERCC spike-in controls to normalize expression relative to rRNA—which remain steady during HSV-1 productive infection— allowing a quantitative comparison of samples with varying levels of host shut off. We quantified differentially expressed viral genes, focusing on viral transcripts upregulated relative to hp66 infection (Fig. 3D). This identified ICP0, ICP47, ICP4, RL1, and LAT as possible candidates responsible for the differential tRNA phenotype in hp66. These regions of the genome also contain viral ncRNA transcripts: Latency-Associated Transcript (LAT), miRNAs, and L/STs. We used genomic lesion mutants (d120, d92, d99, R3616 F-ΔICP47), rather than nonsense mutants, to clarify our findings. Consistent with Fig. 3A-B, we found that the coding regions of ICP0 and ICP4 were not required for tRNA upregulation (Supplemental Fig. 3-1). Additionally, we found mutants with lesions for ICP47, RL1 or LAT induced host tRNA species (Supplemental Fig. 3-1B). Taken together we concluded that ICP0, ICP47, ICP4, RL1, LAT, and miRNAs encoded within LAT or ICP4 were not responsible for tRNA upregulation.

This led us to our final hypothesis, in which the amount of viral transcripts—but not the identity of those expressed—is critical for tRNA upregulation. As expected, we observed robust viral transcription in n212 and wild-type KOS infection (Fig. 3E-F). ICP4, ICP27, and ICP22 all function to promote viral transcript accumulation, thus mutants defective for these proteins had significantly reduced viral transcript levels, ∼7-fold compared to wildtype (Fig. 3D-E). Supporting our hypothesis, hp66 was the most defective for total viral transcription, ∼20-fold compared to wildtype (Fig. 3E-F). These data taken together conclude that tRNA upregulation requires nuclear viral entry, but not synthesis of specific viral transcripts, nascent viral genomes, or viral progeny. These findings support a link between the amount, but not identity, of viral transcripts and tRNA upregulation.

### Changes to Pol III GTF binding after HSV-1 infection

Next, we tested if infection altered recruitment of the Pol III transcription machinery to tRNA loci. We also assessed recruitment of the catalytic subunit of Pol II, POLR2A. We performed ChIP-Seq in MRC5 cells mock-infected or infected with KOS for 2, 4, or 6 hours. Of the transcription machinery tested—POLR3A, BRF1, GTF3C5, TBP, POLR2A—we found that POLR2A had increased binding, by ∼2-fold, to tRNA loci (Fig. 4A-C). Binding of TBP to tRNA loci also showed a moderate increase, but not as strong as POLR2A. Concurrent with prior findings, POLR2A binding to mRNA genes decreased drastically during infection, a phenomenon that promotes host shut off (8-11) (Fig. 4A). tRNA loci with increased POLR2A recruitment were found in accessible regions of the genome (Fig. 4B) and DM-tRNA-Seq results for these loci found increased accumulation after infection (Fig. 4B-C). We did not observe increased recruitment of the catalytic subunit of Pol III, POLR3A, despite our earlier finding that tRNA loci had increased transcriptional output (Fig. 2D-E). The absence of increased Pol III promoter recruitment indicates initiation rates were not solely altered. This suggests that the increased rate of tRNA transcription is caused by a simultaneous increase in Pol III initiation and elongation rates (37). While other studies have found the Pol II and III transcriptional machinery to be linked, it is still unknown exactly how Pol II enhances or regulates Pol III transcription. Further study is required to determine the mechanism by which Pol II enhances tRNA transcription.

**Fig. 4.**
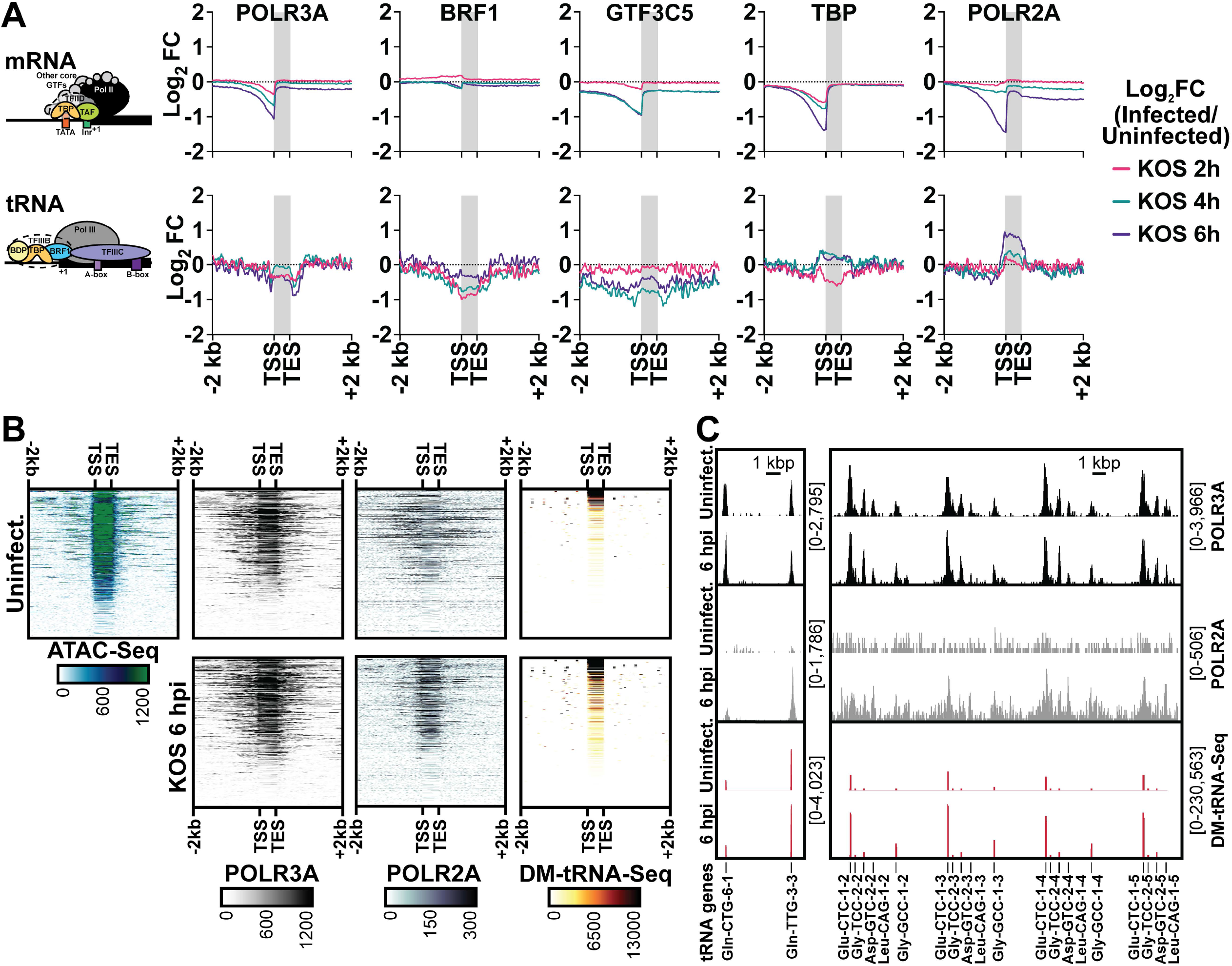
Changes to Pol III GTF binding after HSV-1 infection. A-C) Human fibroblasts were mock-infected or infected with wildtype HSV-1 (KOS) and ChIP-Seq (*A*), ATAC-Seq (*B*), or DM-tRNA-Seq (*C*) was performed. ChIP-Seq data is from biological duplicate experiments and normalized for sequencing depth and cellular genome sampling. ATAC-Seq data is from biological duplicate experiments and normalized for sequencing depth, excluding MT reads. DM-tRNA-Seq data is normalized as described in Fig. 2. A) The average log_2_ fold change for infected over the matched uninfected dataset from 2 kb upstream of TSS to 2 kb downstream of TES for all mRNA or tRNA loci. B) Heatmaps for normalized sequencing data mapped from 2 kb upstream of TSS to 2 kb downstream of TES for tRNA loci. C) Traces of representative tRNA genes, y-axes minimum and maximum are shown in brackets.

To determine whether transcriptional host shut off and tRNA upregulation may be linked phenomena, we performed ChIP-Seq for POLR2A on human fibroblasts after infection with d109, n12, 5dl1.2, n199, and KOS for 6 hours (Fig. S4-1). We observed increased recruitment of POLR2A to tRNA loci for all viruses tested, with the exception of d109 (Fig. S4-1A-B). Host shut off is dependent on the presence of ICP4, and scales with viral genome copy number (10, 11). Consistent with this, we only observed depletion of Pol II from host promoters after infection with 5dl1.2, n199, and KOS infection (Fig. S4-1C-D). The absence of Pol II recruitment to tRNA loci after d109 infection agrees with our earlier findings that d109 infection does not cause tRNA upregulation (Fig. 1 and 3).

To assess how HSV-1 promotes Pol II recruitment at tRNA loci we investigated changes in transcription factor expression and availability. We performed polyA-selected RNA-Seq on human fibroblasts mock-infected or infected with KOS for 2, 4, 6, 8 and 12 h (Fig. S4-2). In line with the global reduction of host mRNA species following HSV-1 infection (Fig. S4-2A), by 12 hpi transcripts for most of the Pol II and III machinery had decreased between 2 to 32-fold (Fig. S4-2B-C). The one exception was POLR2A, for which we observed a slight increase in transcript abundance, peaking around 4 hpi (Fig. S4-2B). We next assessed whether these transcriptional changes resulted in altered protein expression (Fig. S4-3). While POLR2A transcript abundance increased around 4 hpi, its protein expression levels decreased ∼4-fold by 12 hpi (Fig. S4-3B. Again, we observed a marked decrease in protein expression levels for all transcription machinery assessed, with the exception of POU2F1 and SP1. These cellular enhancers are of particular note because they promote Pol III Type III transcription and are also critical for VP16-mediated enhancement of viral immediate early transcription (38-41). Since these enhancers are not required for Pol III type II transcription, we find it unlikely they contribute to tRNA upregulation, however further work is required. Ultimately these results do not explain enhanced Pol II recruitment to tRNA in an environment where their transcription and protein expression levels are globally decreased.

Another mechanism by which HSV-1 alters the host environment is by remodeling the nucleus into sub-domains. HSV-1 assembles within replication compartments enriched for utilized host factors, such as transcription and replication machinery, while excluding negative host factors, most notably histones (42). We imaged Vero cells that were uninfected (Fig. S4-4A), infected pre-replication (Fig. S4-4B), and infected post-replication (Fig. S4-4C-D). We used the nucleoside analog, EdC, to specifically label viral genomes and then stained for various components of the transcriptional machinery including: POLR2A, POLR3A, POLR3G, BRF1, and GTF3C5. We observed a strong reorganization to viral replication compartments for all host factors tested. Of note, only POLR2A colocalized with viral input genomes (Fig. S4-4B). All other components colocalized with the viral genome only after formation of viral replication compartments and multiple rounds of viral genome replication had occurred (Fig. S4-4D). Based on these results, it is unlikely that HSV-1 promotes Pol II recruitment to tRNA loci by enhancing the local concentration gradient. Additionally, most factors reorganized to viral replication compartments (Fig. S4-4), suggesting decreased recruitment to the host genome—an observation in line with our ChIP-Seq results showing global decrease of POLR2A and TBP from host mRNA promoters and slight decrease of POL3A, BRF1, and GTF3C5 from tRNA loci (Fig. 4A). These results mirror the global environment of host transcriptional repression present in HSV-1 infection, wherein tRNA appear to be the rare outlier.

### Recruitment of Pol III to the HSV-1 genome

Classically all HSV-1 transcripts are defined as Pol II-dependent (43). However, there are examples of other DNA viruses that encode Pol III transcripts; the VAI and VAII RNAs of adenovirus (14), EBER1 and EBER2 of Epstein Barr Virus (15), and the tRNA-micro RNA (miRNA) encoding RNAs (TMERs) of MHV68 (16-18). We examined our ChIP-Seq data for binding of the Pol III machinery to the HSV-1 genome. We observed strong, distinct recruitment of the catalytic subunit of Pol III, POLR3A, to the viral genome (Fig. 5A). Of the transcription factors tested—POLR3A, POLR2A, TBP, GTF3A, GTF3C5, BRF1, BRF2, ICP4—POLR3A binding most closely resembled POLR2A (Fig. 5B). POLR3A binding to the viral genome was most prominent during early viral infection (pre-replication), and occurred independent of the major viral transcriptional activator, ICP4 (Fig. S5-1 and S5-2). There were a few prominent instances where POLR3A binding did not mimic POLR2A, namely within the latency associated transcript (LAT) intron, at the promoter of RL1, and intergenic between US9 and US10.

**Fig. 5.**
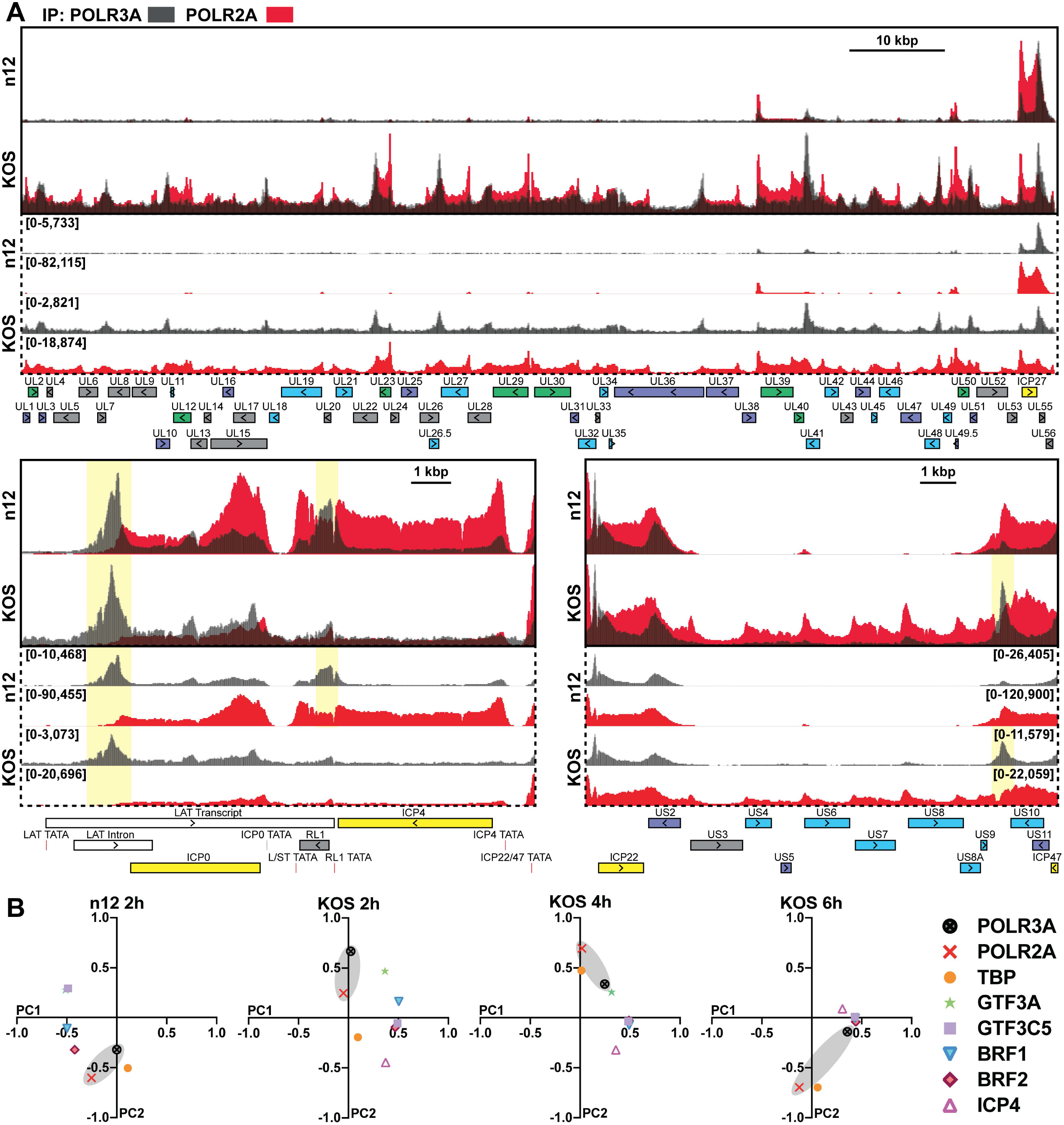
Recruitment of Pol III to the HSV-1 genome. Human fibroblasts were infected with n12 (ΔICP4) or KOS (WT) HSV-1 for 2 h. ChIP-Seq was performed for POLR3A, POLR2A, TBP, GTF3A, GTF3C5, BRF1, BRF2, and ICP4. Data is the average of biological duplicates and normalized for sequencing depth and viral genome copy number. A) Traces of POLR2A and POLR3A binding to the unique long (UL), joint, and unique short (US) regions of the HSV-1 genome (KT899744.1 assembly). Y-axes maximum and minimum values are listed within brackets. Viral CDS are listed below, with colors indicating transcriptional class: immediate early (yellow), early (green), leaky late (blue), true late (purple), unclassified (grey). We have also annotated additional genomic features such as TATA boxes (red) and ncRNA (white). Regions where POLR3A bound distinct from POLR2A are highlighted in yellow. B) Principal component analysis of GTF binding profiles for the viral genome, these were calculated by binning (10 bp) normalized traces.

### Unique GTF context of viral Pol III binding

A closer analysis of the Pol III machinery recruited to the viral genome revealed two POLR3A binding contexts: 1. Pol II Coincident—with TBP, GTF3C5, GTF3C6, and GTF3A, or 2. Pol II Independent—with GTF3C1 and BRF2 (Fig. 6). These binding contexts are drastically different from those characterized (19, 20) and observed in our own data for host promoters (Fig S6-1 and S6-2). Notably the components of the TFIIIC complex, GTF3C1 to GTF3C6, always bind coincident on host Pol III type II promoters. On the HSV-1 genome, binding of GTF3C2 and GTF3C3 was largely absent (Fig. S6-3). GTF3C1, GTF3C5, and GTF3C6 were recruited to the viral genome, however the binding pattern of GTF3C1 did not coincide with the other TFIIIC subunits present (Fig. S6-3). Perhaps the most interesting binding pattern is for the Pol III Type III transcription factor, BRF2. This factor has very select host binding with around a dozen targets (19), an observation consistent with our dataset (Fig. S6-1). BRF2 bound robustly within the LAT intron, coincident with a Pol II-independent POLR3A binding event (Fig. 6). This binding context requires further investigation to identify what viral transcripts may be produced from this locus.

**Fig. 6.**
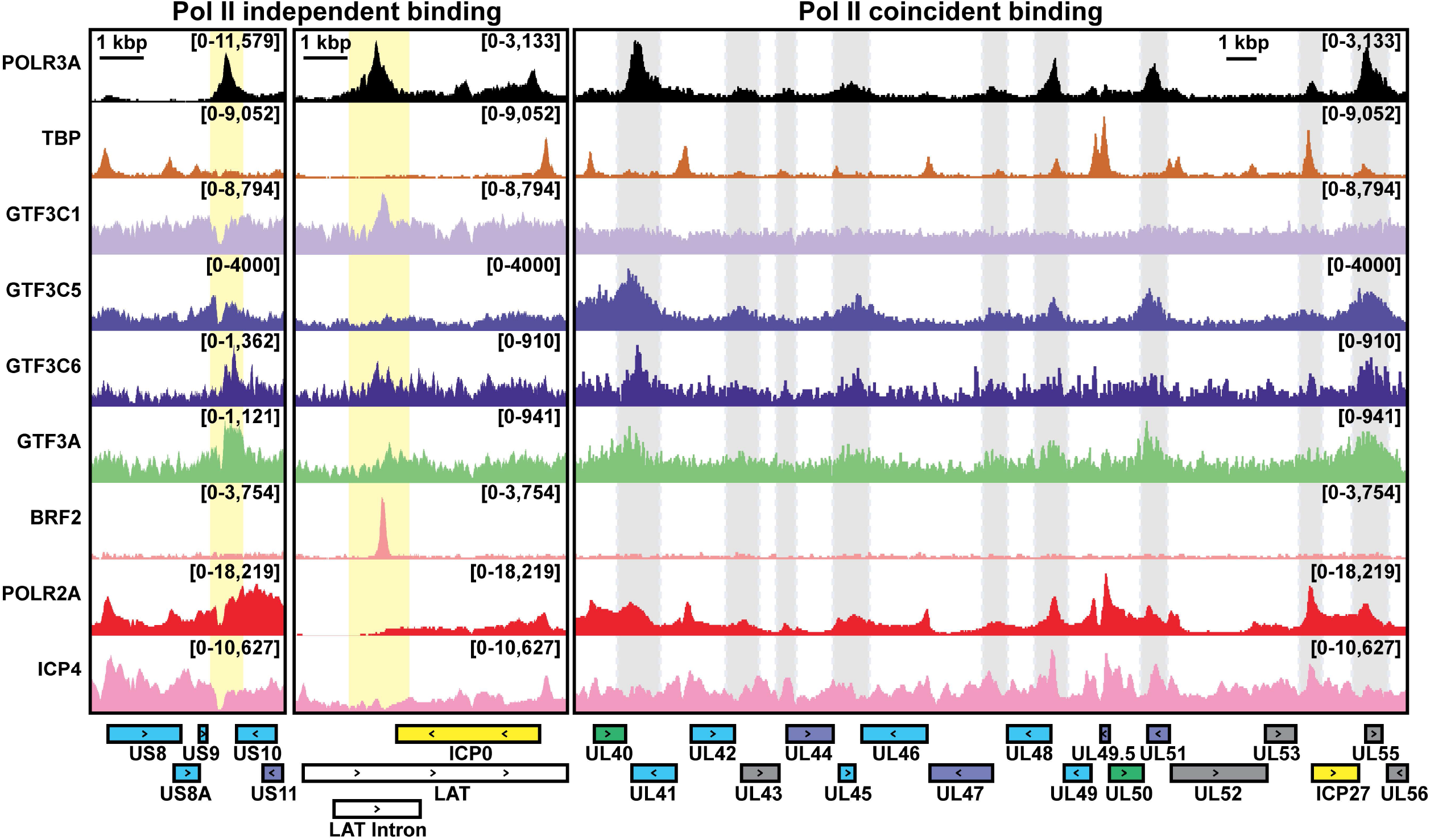
Unique GTF context of viral Pol III binding. Human fibroblasts were infected with KOS (WT) HSV-1 for 2 h. ChIP-Seq was performed for POLR3A, TBP, GTF3C1, GTF3C5, GTF3C6, GTF3A, BRF2, POLR2A, and ICP4. Data for GTF3C2, GTF3C3, GTF3C4 is not shown, as we did not observe recruitment to the viral genome (Fig. 6-1). Data is normalized and presented as in Fig. 5. Yellow or grey boxes were used to highlight regions of Pol II-independent or – coincident binding, respectively.

## DISCUSSION

Herein we characterized how HSV-1 affects Pol III mediated transcription (Fig. 7), a pathway commonly dysregulated by cancer and pathogens. In prior work, HSV-1 has been shown to induce the Pol III type II transcript, Alu (30, 31). Echoing these results, we found that HSV-1 targets and upregulates additional Pol III type II transcripts. This includes a 2-fold increase in total abundance of tRNA, and a 10-fold increase in nascent levels of tRNA. We did not observe increased recruitment of Pol III to tRNA. Given these results, we propose that tRNA upregulation is caused by a simultaneous increase in Pol III initiation and elongation rates (37). Given the short half-life of pre-tRNAs, decreased turnover may also contribute to increased nascent abundance. Further work would need to be done to determine whether turnover rates are altered during HSV-1 infection.

**Fig. 7.**
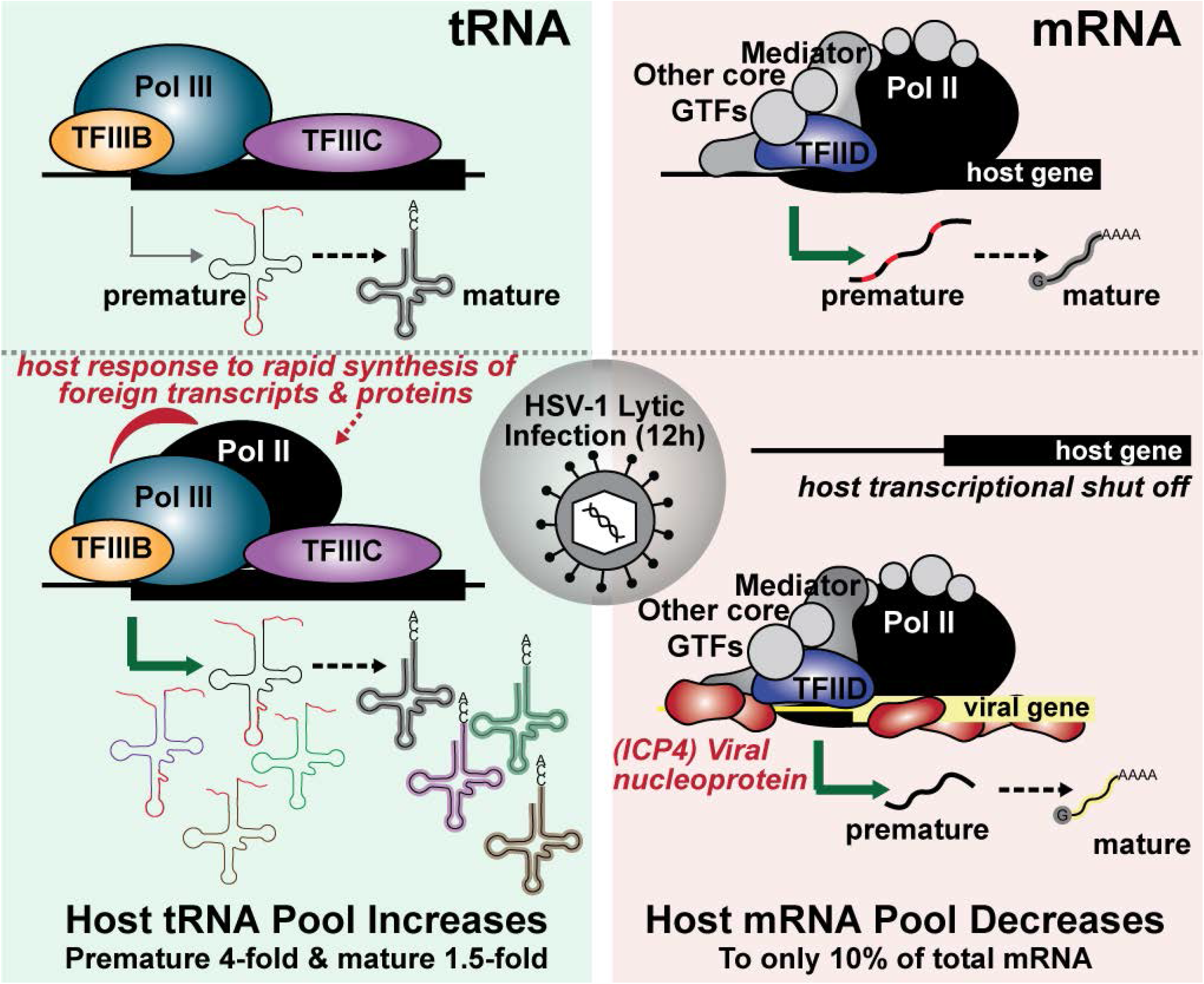
Model of HSV-1 Pol II and III Modulation during lytic infection. During lytic infection, HSV-1 alters host Pol II recruitment and induces depletion from host mRNA in favor of host tRNA and viral mRNA. With 12 hours, these changes result in a 2-fold increase in total abundance of tRNA and a 10-fold increase in nascent levels of tRNA. In contrast, host mRNA decreases to only 10% of the total present in the cell. These phenomena are mediated distinctly. Depletion of Pol II from mRNA promoters scales with viral genome replication and ICP4. Recruitment of Pol II to tRNA promoters, and their subsequent upregulation, is caused by entry of the viral genome and synthesis of viral transcripts—with no dependence on exactly which transcripts are made.

We identified 15 mature-and 66 pre-tRNA upregulated within 12 hours of HSV-1 infection. Exactly which, if any, of these upregulated tRNA species may be rate limiting during viral expression requires further study. There is evidence among other viruses for this hypothesis, for instance HIV lytic replication induces tRNA expression (44, 45). This is so critical that the host innate immune response combats HIV by targeting this pathway via the interferon responsive gene, SLFN11 (46). HSV-1 mediated tRNA upregulation is at odds with the general environment of host transcriptional shut off mediated by the virus, including downregulation of most protein-coding mRNA and Pol III type III transcripts. We posit that HSV-mediated tRNA upregulation is required for robust productive replication, wherein a single infected cell can synthesize up to 30,000 viral progeny within a 12-hour window. During this time the virus must produce ∼90 viral proteins at exponential rates to facilitate genome replication and virion assembly. Alternatively, increased tRNA expression may be critical in infection scenarios where host shut off is less pronounced and there is more competition for translation resources.

Another possible option is that an increase in Pol III transcripts may result in an overabundance of RNA molecules that have exposed 5’-triphosphates, which serve as substrates for RNA sensors, such as RIG-I (47). HSV-1 effectively blocks the innate immune response during productive infection—the exact mechanism behind nuclear sensing is still at question (48). HSV-1 perturbation of the Pol III landscape may have a negative side effect in stimulating an innate immune pathway that the virus must quickly block or outstrip in a bid to productively replicate.

To determine the mechanism by which HSV-1 induces tRNA transcription, we tested various hypotheses including altered Pol III transcription factor expression, recruitment, or localized concentration. HSV-1 productive infection resulted in a decrease for all three. Decreased Pol III machinery expression suggests that the mechanism by which HSV induces tRNA synthesis differs from other viruses. Namely adenovirus (23-25), SV40 (26), and Epstein Barr Virus (27) which increase the abundance of Pol III transcription factors by viral regulatory proteins E1A, E1B, T-antigen, and EBNA1. We then turned to another possible option, namely crosstalk between the Pol II and III transcription machinery. Recent work has increasingly demonstrated a dependence and interplay between the Pol II and III machinery (19-22, 49). In a global environment of Pol II depletion from host genes, we were surprised by a 2-fold increase in Pol II recruitment to tRNA loci. Recruitment of Pol II to tRNA genes only occurred upon infection of HSV-1 mutants that also had increased tRNA abundance. Additionally, recruitment of Pol II to tRNA genes occurred independent of host transcriptional shut down. The question now becomes, how is Pol II enhancing tRNA transcription rates? We propose three potential options: i. Pol II alters the chromatic environment at bound tRNA, ii. enhances the rate of Pol III termination or re-initiation, or iii. functions itself to transcribe tRNA.

Infection with a panel of HSV-1 mutants allowed us to closely characterize viral life events critical for tRNA upregulation. We found that nuclear viral entry was required, but synthesis of E and L transcripts, nascent viral genomes, and viral genomes was not required. Furthermore host shut off was not necessary for tRNA upregulation, suggesting a different mechanism of induction than MHV68 perturbation of the tRNA landscape (29). Our results define a narrow window of viral processes which may be critical for tRNA upregulation, namely entry of the viral genome and synthesis of viral transcripts—with no dependence on exactly which transcripts are made. We propose that changes in the pool of tRNA may be caused by a common feature among HSV-1 transcripts, including: GC content of ∼68%, propensity to form G-quadruplexes, and high density of complementary transcripts. Alternatively, tRNA abundance profiles are known to change drastically in response to various stimuli including oxidative stress, osmotic stress, temperature stress, and diauxic shift (50). Productive viral infection takes an extreme toll on host cell homeostasis, altering levels of metabolites including nucleotides and amino acids (51), inducing an unfolded protein response (52), and dysregulating the DNA damage response (53). Most notably HSV-1 transcripts are composed from a high density of intrinsically disordered domains, and due to the rapid rate of synthesis are prone to misfolding and aggregation—traits that robustly trigger a UPR. Any one of these cellular feedback loops could be hijacked by HSV-1 to increase the pool of tRNA immediately available for viral use.

Thus far, we have focused on how the virus alters host Pol III transcription. However, various viruses including adenovirus, Epstein Barr Virus, and murine herpesvirus-68 possess Pol III-dependent viral transcripts (14-18). These RNAs have functions that combat the innate antiviral response, contribute to latency, and play a role in transformation. Our data suggests that Pol III may play novel roles in HSV-1 transcription. We observed Pol III binding to the viral genome in two conformations: i. coincident with RNA Pol II, or ii. coincident with the canonical Pol III type III transcription factor, BRF2. The latter binding event occurred in a region of the genome where latency-derived transcripts originate, suggesting a cell-type specificity and potential role in viral persistence and reactivation. These L/S joint region of the viral genome is rich with ncRNA, including LAT (54), L/STs (55), and microRNAs (56). Pol II promoters have been associated with the genesis of the LAT and L/STs (57, 58), and we have also shown that these promoters function in a reconstituted Pol II in vitro transcription systems (59, 60). However, these studies do not preclude the possibility that there are alternative mechanisms to transcribe these RNAs, the microRNAs, or other noncoding RNAs yet to be discovered. Herein we provide the first evidence for a putative Pol III transcript derived from the HSV-1 genome.

## ACKNOWLEDGEMENTS

**Thanks to Hannah Fox for datasets and Jill Dembowski for thoughtful discussions. NIH grants R01-AI030612 and R21-AI156065 to N.A. D. NIH grants T32-AI060525 and F31-AI36251 to S. E. D. American Cancer Society Postdoctoral Award 131370-PF-17-245-01-MPC to J.M.T. NIH grant R01-AI147183 to B.A.G.**

## FIGURE LEGENDS

**Supplemental Table 1 DM-tRNA-Seq data**

Human fibroblasts were mock-infected or infected with ΔICP4 (n12) or wildtype HSV-1 (KOS) for 12 hours. DM-tRNA-Seq was performed, data is the average of four data points consisting of two biological replicate experiments each containing two technical replicates. Data was normalized to an internal spike-in control and the size in kb of each tRNA.

**Supplemental Fig. 2-1 Distribution of tRNA isodecoders changed in HSV-1 infection.**

Analysis of isodecoder frequency among DE pre-and mature-tRNA from the DM-tRNA-Seq dataset. “DE-Up” had Log_2_ fold change of KOS/uninfected of ≥ 0.5 and p-value of < 0.05. “DE-Down” had Log_2_ fold change of KOS/uninfected of ≤ −0.5 and p-value of < 0.05. As tRNA genes are degenerate we summarized by target and anticodon (isodecoder). A) Distribution of affected tRNA isodecoders as a function of DE targets. B) Using the annotated coding sequence of HSV-1 we broke down the theoretical distribution of codon usage, assuming all viral proteins are made equally. Since the HSV-1 genome is ∼68% GC content, isodecoder usage is skewed towards those with GC-rich codons. C) Distribution of affected tRNA isodecoders as a function of those expressed in our experimental system. To classify a tRNA as expressed or “detected” in our experimental system, we required a cut off of at least 100 normalized mapped reads in uninfected or infected conditions.

**Supplemental Fig. 2-2 Relative localization of DE tRNA genes to other host genes.**

Analysis of relative genomic position among DE pre-and mature-tRNA from the DM-tRNA-Seq dataset. “DE-Up” had Log_2_ fold change of KOS/uninfected of ≥ 1 and p-value of < 0.05. “DE-Down” had Log_2_ fold change of KOS/uninfected of ≤ −1 and p-value of < 0.05. A-B) Relative position was analyzed for DE tRNA to promoters from all annotated genes, excluding tRNA. C-D) Relative position was analyzed for DE tRNA to promoters from only protein-coding genes. A, C) Relative position as a function of all tRNA analyzed. B, D) Percentage of total targets that were located with 3 kb of a promoter.

**Supplemental Fig. 3-1 Refining HSV-1 gene expression required for tRNA upregulation.**

A, D) Details of strain genotypic and phenotype differences. B) Human fibroblast cells were mock-infected or infected with n12, d120, d92, 5dl1.2, n212, d99, n199, R3616, wild-type HSV-1 strain KOS, or wild-type HSV-1 strain F. RNA was isolated at 12 hpi, and Northern blots were used to assess transcript abundance. Representative images from biological duplicates are shown. C) 300 ug/mL phosphonoacetic acid (PAA) or 100 uM acyclovir (ACV) was used to treat mock-infected or KOS-infected cells at 0 h. RNA was collected at 12 h and assessed by Northern blots. Representative images from biological duplicates are shown.

**Supplemental Fig. 4-1 Pol II depletion from host mRNA and enrichment on tRNA are not linked.**

Human fibroblasts were mock-infected or infected with d109 (ΔICP0/4/22/27/47), n12 (ΔICP4), 5dl1.2 (ΔICP27), n199 (ΔICP22), or wild-type HSV-1 (KOS) and ChIP-Seq for POLR2A was performed. ChIP-Seq data is from biological triplicate experiments and normalized for sequencing depth and cellular genome sampling. The average log_2_ fold change of infected/uninfected signal from 2 kb upstream of TSS to 2 kb downstream of TES for all mRNA or tRNA loci. A,C) Data plotted is average of biological triplicates (solid line) with standard deviation (dashed line, shaded). B, D) Data plotted is average of biological triplicates.

**Supplemental Fig. 4-2 Transcriptomic changes of Pol II and III machinery during HSV-1 productive infection**

Human fibroblasts were mock-infected or infected with wildtype HSV-1 (KOS) for the indicated times. PolyA-selected RNA-Seq was performed. Data is the average of biological triplicate experiments and error bars are standard deviation. A) Global changes are represented as number of reads mapped to viral or host genome per total reads sequenced (% Total Reads). B-C) Mapped reads per billion total reads per kilobase pair (MR/BTR/Kb), or log_2_ fold change of infected over uninfected cells (Log_2_FC).

**Supplemental Fig. 4-3 Proteomic changes of Pol II and III machinery during HSV-1 productive infection**

A) Tandem mass tag mass spectrometry (TMT-MS) data from mock-or HSV-1 infected HaCaT cells (human immortalized keratinocytes), published in (61). B) Human fibroblast (MRC5) cells were mock-infected or infected with d109 (ΔIE’s) for 4 hours, n12 (ΔICP4) for 2 hours, or WT (KOS) HSV-1 for 2, 4, 6, 8 or 12 hours. Protein was isolated and measured via LiCor-Western blot for indicated proteins. Each data point is a biological replicate.

**Supplemental Fig. 4-4 Altered nuclear organization of the Pol III transcription machinery**

Vero cells were A) mock-infected or B-D) infected with wildtype HSV-1 (KOS) for the indicated time. B) To image input viral genomes we infected with an EdC-prelabed viral stock. C-D) To image nascently replicated viral genomes we pulsed EdC at the indicated time points before fixing. Cells were fixed and EdC labeled DNA was tagged with alexa fluor 488 to visualize viral genomes (green) and cellular proteins were visualized by immunofluorescence (red). Nuclei were labeled with Hoechst (blue). Images are representative of biological duplicate experiments.

**Supplemental Fig. 5-1 Pol III machinery recruitment to the HSV-1 genome during early infection**

Human fibroblasts were infected with n12 (ΔICP4) or KOS (WT) HSV-1 for 2 h. ChIP-Seq was performed for POLR3A, POLR2A, TBP, GTF3A, GTF3C5, BRF1, and BRF2. Data is the average of biological duplicates and normalized for sequencing depth and viral genome copy number. Traces of binding to the unique long (UL), joint, and unique short (US) regions of the HSV-1 genome (KT899744.1 assembly). Y-axes maximum and minimum values are listed within brackets. Viral CDS are listed below, with colors indicating transcriptional class: immediate early (yellow), early (green), leaky late (blue), true late (purple), unclassified (grey). We have also annotated additional genomic features such as TATA boxes and ncRNA (white).

**Supplemental Fig. 5-2 Pol III machinery recruitment to the HSV-1 genome post-viral genome replication**

Human fibroblasts were infected with KOS (WT) HSV-1 for 4 or 6h. ChIP-Seq was performed for POLR3A, POLR2A, TBP, GTF3A, GTF3C5, BRF1, and BRF2. Data is the average of biological duplicates and normalized for sequencing depth and viral genome copy number. Traces of binding to the unique long (UL), joint, and unique short (US) regions of the HSV-1 genome (KT899744.1 assembly). Y-axes maximum and minimum values are listed within brackets. Viral CDS are listed below, with colors indicating transcriptional class: immediate early (yellow), early (green), leaky late (blue), true late (purple), unclassified (grey). We have also annotated additional genomic features such as TATA boxes and ncRNA (white).

**Supplemental Fig. 6-1 Binding of Pol III machinery to canonical cellular targets.**

ATAC-Seq or ChIP-Seq for POLR3A, TBP, BRF1, GTF3C5, GTF3A, BRF2, and POLR2A was performed on uninfected human fibroblasts. Heatmaps are from 2 kb upstream of TSS to 2 kb downstream of TES for loci, and “n” indicates the number of loci contained within each heatmap. y-axes are mapped reads per billion total reads (MR/BTR). Models of canonical Pol III promoter architecture was adapted from (19).

**Supplemental Fig. 6-2 TFIIIC complex recruitment to canonical cellular targets.**

ATAC-Seq or ChIP-Seq for GTF3C1, GTF3C2, GTF3C3, GTF3C4, GTF3C5, and GTF3C6 was performed on uninfected human fibroblasts. Heatmaps are from 2 kb upstream of TSS to 2 kb downstream of TES for loci, and “n” indicates the number of loci contained within each heatmap. y-axes are mapped reads per billion total reads (MR/BTR). Models of canonical Pol III promoter architecture was adapted from (19).

**Supplemental Fig. 6-3 TFIIIC complex recruitment to the HSV-1 genome during early infection**

Human fibroblasts were infected with KOS (WT) HSV-1 for 2 h. ChIP-Seq was performed for POLR3A, POLR2A, GTF3C1, GTF3C2, GTF3C3, GTF3C4, GTF3C5, and GTF3C6. Data is the average of biological duplicates and normalized for sequencing depth and viral genome copy number. Traces of binding to the unique long (UL), joint, and unique short (US) regions of the HSV-1 genome (KT899744.1 assembly). Y-axes maximum and minimum values are listed within brackets. Viral CDS are listed below, with colors indicating transcriptional class: immediate early (yellow), early (green), leaky late (blue), true late (purple), unclassified (grey). We have also annotated additional genomic features such as TATA boxes and ncRNA (white).

## METHODS

### Cells and viruses

Vero (African green monkey kidney), U2OS (human osteosarcoma), and MRC5 (human fetal lung) cells were obtained from and propagated as recommended by ATCC. Viruses used in this study include HSV-1 mutants: n199 (ICP22 null, (62)), n212 (ICP0 null, (63)), d99 (ICP0 null, ((64))), n12 (ICP4 null, (65)), d120 (ICP4 null, (34)), 5dl1.2 (ICP27 null, (66)), d92 (ICP4/27 null, (67)), d109 (ICP0/ICP4/ICP22/ICP27/ICP47 null, (64)), R3616 (RL1 null, (68)), F-ΔICP47 (ICP47 null, (69)), F-ΔICP47-R (ICP47 null revertant, (69)), hp66 (UL30 null, (70)), and wild type HSV-1 (KOS, (71)). n199, R3616, F-ΔICP47, and KOS were prepared and titered in Vero cells. Other virus stocks were prepared and titered in the following Vero-based complementing cell lines: E5 (ICP4+, n12, d120-complementing, (72)); E11 (ICP4/ICP27+, 5dl1.2, d92-complementing, (67)); F06 (ICP4/ICP27/ICP0+, d109-complementing, (73)); U2OS (n212, d99-complementing), POLB3 (hp66-complementing, (70)). We thank Don Coen (hp66, POLB3), David A. Leib (R3616), and Anthony St. Leger (F-ΔICP47, F-ΔICP47-R) for their kind gift of viruses or cells.

### Antibodies

A list of all antibodies and the amount used per technique is located in Table 1.

**Table 1.** Antibodies Used in Study.

| Target | Antibody | Amount/Dilution Used |  |  |
| --- | --- | --- | --- | --- |
|  |  | ChIP-Seq | Immunofluorescence | Western Blot |
| α-Tubulin | AbCam #ab7291 | N/A | N/A | 1:5000 |
| Vinculin | Abcam #ab129002 | N/A | N/A | 1:15000 |
| GAPDH | ThermoFisher #AM4300 | N/A | N/A | 1:5000 |
| POLR2A | AbCam #ab5408 | 5 μg/IP | 1:500 | N/A |
| POLR2A | SantaCruz #sc899 | N/A | N/A | 1:500 |
| TBP | AbCam #ab51841 | 25 μg/IP | N/A | 1:1500 |
| POLR3A | AbCam #ab96328 | 10 μg/IP | 1:300 | N/A |
| POLR3A | CST #12825S | N/A | N/A | 1:500 |
| POLR3B | AbCam #ab137030 | N/A | N/A | 1:250 |
| POLR3C | Bethyl #A303-064A-M | N/A | N/A | 1:250 |
| POLR3D | AbCam #ab86786 | N/A | N/A | 1:500 |
| POLR3E | Sigma #HPA041477 | N/A | N/A | 1:125 |
| POLR3F | Abcam #ab180501 | N/A | N/A | 1:500 |
| BRF1 | SantaCruz #sc-390821 | 10 μg/IP | 1:200 | 1:100 |
| BRF2 | SantaCruz #sc-390312 | 10 μg/IP | N/A | 1:100 |
| GTF3A | Abcam #ab254632 | 1 μg/IP | N/A | 1:250 |
| GTF3C1 | Novus Biologicals #NB100-60657 | 10 μg/IP | N/A | 1:500 |
| GTF3C2 | AbCam #ab89113 | 7.5 μg/IP | N/A | 1:100 |
| GTF3C3 | SantaCruz #sc-101176 | 7.5 μg/IP | N/A | 1:100 |
| GTF3C4 | Sigma #HPA069369 | 3 μg/IP | N/A | 1:500 |
| GTF3C5 | Bethyl #A301-242A | 3 μg/IP | 1:500 | 1:500 |
| GTF3C6 | ThermoFisher #PA5-63948 | 7.5 μg/IP | N/A | 1:100 |

### Oligos

A list of all single stranded oligos used for either qPCR or Northern Blot is located in Table 2.

**Table 2.**
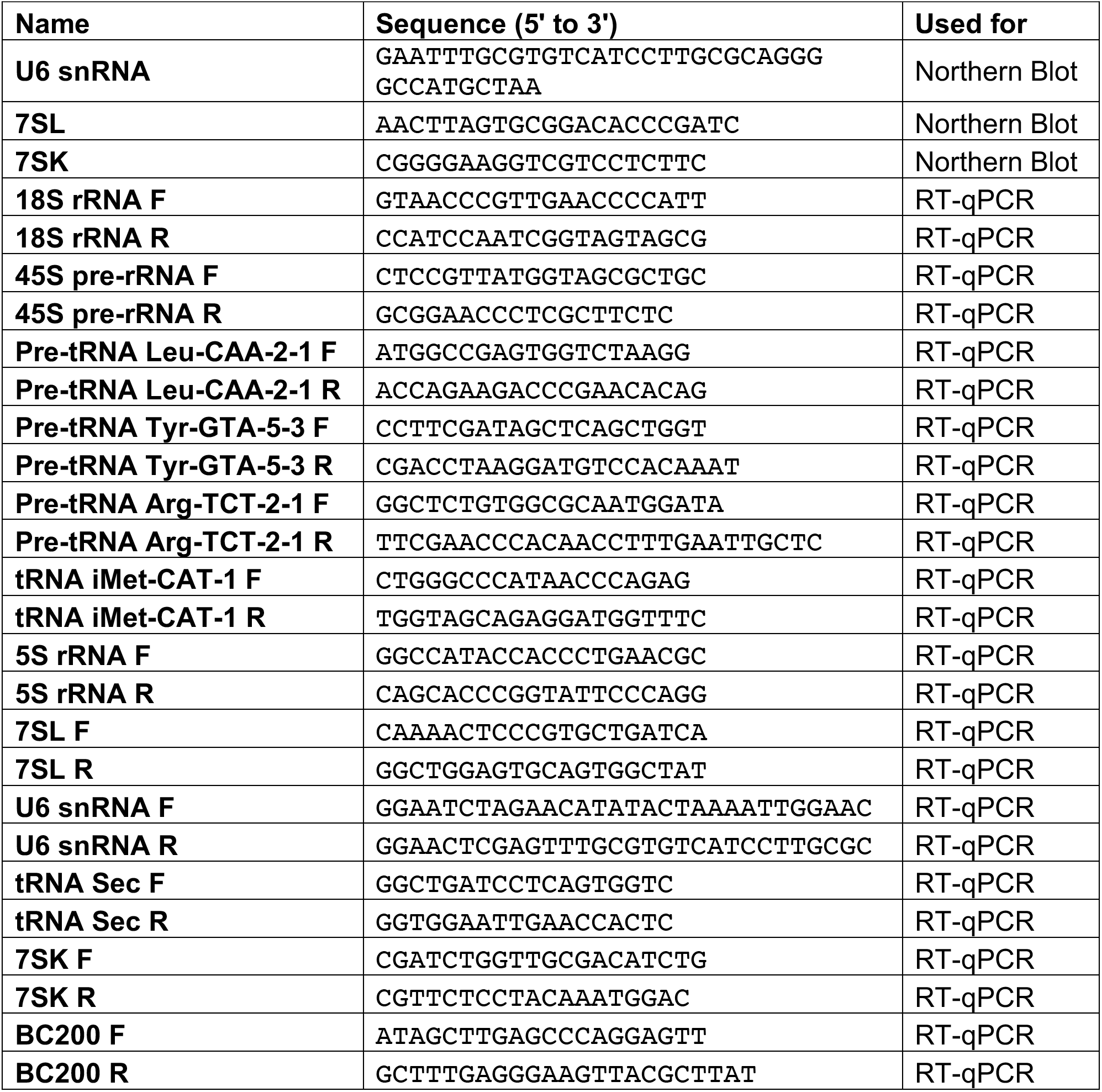
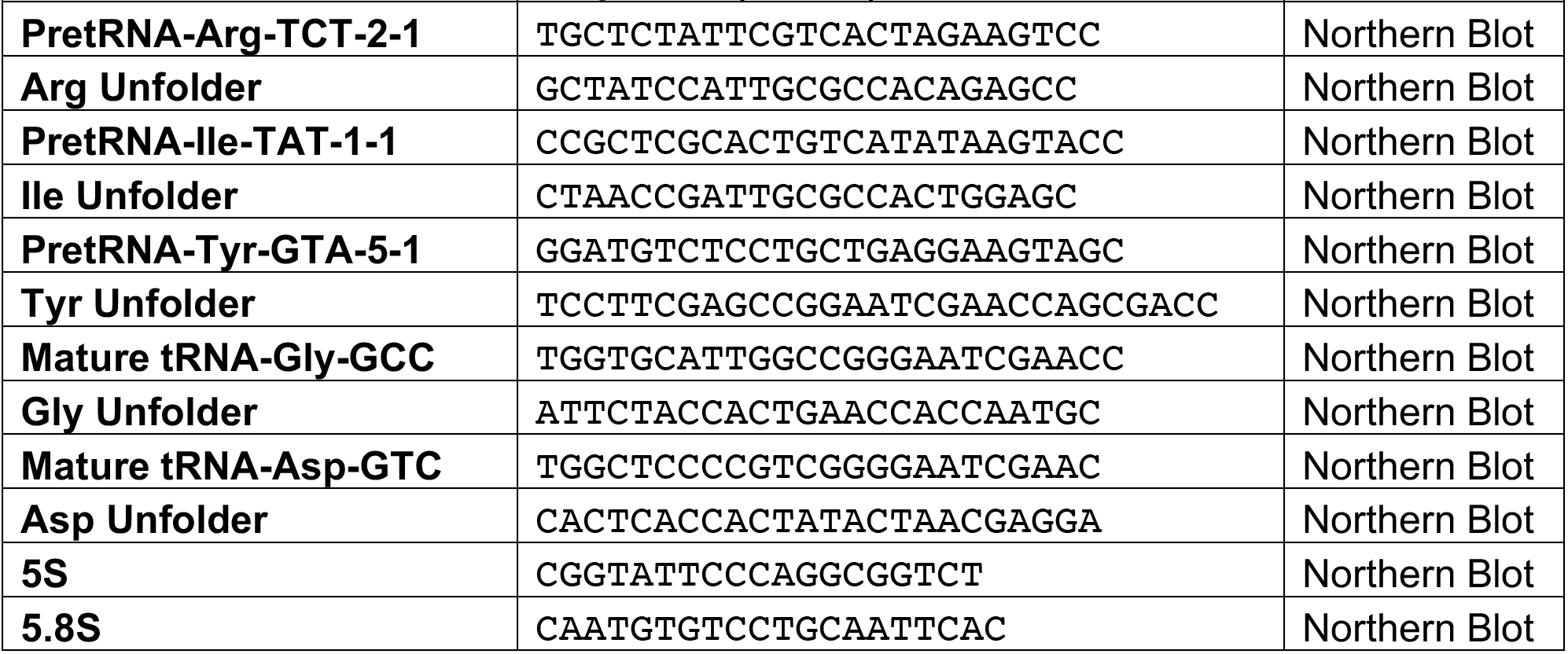
Oligos Used in Study.

### Viral infection

Confluent cell monolayers were infected with 10 PFU per cell. Virus was adsorbed in tricine-buffered saline (TBS) for 1 hr at room temperature. Viral inoculum was removed, and cells were washed quickly with TBS before adding 2% FBS media. 0 hour time point was considered after adsorption of infected monolayers when cells were place at 37°C to incubate.

### qPCR Quantification of mRNA

Cell monolayers were collected by aspirating supernatant and washing twice with TBS. Cells were scraped into 1 mL TBS and pelleted, supernatant was discarded. RNA was isolated using the RNAqueous Micro Kit (ThermoFisher cat. no. AM1931). cDNA was generated from 500 ng total RNA, as quantified using the Agilent RNA 6000 Nano Kit. RNA was reverse transcribed with 20 units Riboguard RNase inhibitor, 2.5 uM Random decamer primer (Invitrogen cat no. 5722G), 100 4 units MMLV-HP reverse transcriptase, 10 mM dithiothreitol, 2.5 mM dNTPs, and 1x reaction buffer (Epicentre cat no. RT80125K and RG90910K). RNA and random decamer primer were first incubated at 85°C for 3 minutes and then incubated on ice for 2 minutes during which remaining reaction components were added. The entire reaction was incubated at 65°C for 2 minutes and then 37°C for 1 hour. To heat inactivate components the cDNA was incubated at 85°C for 5 minutes. Standard curves were generated using purified KOS or human genome stocks.

### Western Blot

At the indicated times postinfection, proteins were isolated from cells using Laemmli SDS sample buffer and Western blotting was carried out using the primary antibodies listed in table 1. Blot were probed with secondary antibodies—IRDye goat anti rabbit or goat anti-mouse 680/800—at a 1:15000 dilution. The intensity of gel bands was quantified using the Odyssey CLx system machine and Image Studio program. Band intensities were normalized to loading controls including GAPDH, alpha-tubulin (TUB4A), or Vinculin (VCL) detected from the same sample.

### Northern Blot

Total RNA was isolated from cells using TRIzol (Invitrogen), following manufacturer’s instructions. RNA was quantified using Agilent RNA 6000 Nano kit. 5-20 μg of RNA was loaded into 10% TBE-Urea Polyacrylamide Gel, transferred to Hybond-N+ (Sigma) membranes using the Owl™ HEP Series Semidry Electroblotting Systems (ThermoFisher). Blots were pre-hybridized in ExpressHyb buffer (Takara Bio) at 37°C for 1 hour. If using radiolabeled probes specific to tRNAs, 50 nM unfolder oligo (see Table 2) was included in the pre-hybridization step, as described in (74). Probes were generated by end-labeling oligos listed in Table 2 using T4 PNK and [g-32P]-ATP. Probes were hybridized to the blots in ExpressHyb buffer at 37°C overnight. If using radiolabeled probes specific to tRNAs, 25 nM unfolder oligo was included in the hybridization step. After hybridization, blots were washed four time in 2x SSC [0.3M NaCl, 0.03M Trisodium Citrate pH 7.0], 0.05% SDS and 2 times in 0.1x SSC, 0.1% SDS. Blots were exposed and quantified using the Typhoon Biomolecular Imagine (Amersham). Blots were stripped in 1% SDS, 0.1x SSC, 40 mM Tris pH 8 for four 10 minute washes at 70°C. Northern blot quantifications were normalized to 5.8S as a loading control.

### Immunofluorescence

1.7 x 10^5^ Vero cells were infected as described above. Coverslips were fixed at indicated time point with 3.7% paraformaldehyde. EdC labeling of viral replication compartments, click chemistry, and immunofluorescence were conducted as previously described (42, 75). Cellular DNA was stained with 1:2000 Hoescht, and immunofluorescence was carried out using antibodies listed in table 1 and 594-conjugated secondary antibodies (Santa Cruz, 1:500). Images were taken using an Olympus Fluoview FV1000 confocal microscope. Images are representative of biological duplicate experiments.

### DM-tRNA-Seq

Protocol was modified from (29, 35), with the following modifications. 7 x 10^7^ MRC5 cells were infected as described above and total RNA was isolated from cells using TRIzol (Invitrogen), following manufacturer’s instructions. Total RNA extracted from two biological replicates was spiked with *in vitro* transcribed *E. coli* tRNA-Lys, *E. coli* tRNA-Tyr, and *S. cerevisiae* tRNA-Phe transcripts at 0.01 pmol IVT tRNAs per μg total RNA. RNA was deacylated in 0.1M Tris-HCl, pH 9 at 37 °C for 45 min, ethanol purified, and then dephosphorylated with PNK. Deacylated and dephosphorylated RNAs were purified with a mirVANA small RNA purification kit (Ambion). RNAs were demethylated in 300 mM NaCl, 50 mM MES pH 5, 2 mM MgCl2, 50 μM ferrous ammonium sulfate, 300 μM 2-ketoglutarate, 2 mM L-ascorbic acid, 50 μg/ml BSA, 1U/μl SUPERasin, 2X molar ratio of wt AlkB, and 4X molar ratio of D135S AlkB for 2 hours at room temperature. Ni-NTA cation exchange purified His-tagged wild-type and D135S AlkB. Reaction was quenched with 5 mM EDTA and purified with Trizol LS reagent. Two TGIRT reactions from each biological replicate was performed, these are considered technical replicates. In total each sample condition is an average of 4 data points, consisting of two biological replicates containing technical duplicates. 100 ng demethylated small RNAs was used for library prep with a TGIRT Improved Modular Template-Switching RNA-seq Kit (InGex) following the manufacturer’s instructions. PCR amplification was performed with Phusion polymerase (Thermo Fisher) with Illumina multiplex and barcoded primers. Libraries were quantified using the Agilent DNA 7500 Kit, and samples were mixed together at equimolar concentration. Samples were size selected for 150-250 bp fragments using the Pippin system. Illumina NextSeq 550 platform was used to generate 75 bp PE reads and carried out at the Tufts University Core Facility.

### 4SU-Sequencing

2×10^6^ MRC5 cells were infected with wild-type HSV-1 (KOS) as described above. At indicated time (hpi) 500 uM 4SU (Sigma) was added to cell culture medium. 15 minutes post-4SU addition, cells were collected and RNA extracted using miRNeasy kit (Qiagen). 50-100 ug total RNA was biotinylated in 10 mM Tris pH 7.4, 1 mM EDTA, 0.2 mg/mL EZ-link Biotin-HPDP (ThermoFisher). Unbound biotin was removed by performing a chloroform:isoamyl alcohol extraction using MaXtract High Density tubes (Qiagen). RNA was isopropanol precipitated and resuspended in water. Biotinylated RNA was bound 1:1 to Dynabeads My One Streptavidin T1 equilibrated in 10 mM Tris pH 7.5, 1 mM EDTA, 2 M NaCl. Bound beads were washed three times with 5 mM Tris pH 7.5, 1 mM EDTA, 1 M NaCl. 4SU-RNA was eluted with 100 mM DTT and isolated using the RNeasy MinElute Cleanup Kit (Qiagen). RNA-Seq libraries were generated using NEBNext Ultra Directional RNA Library Prep Kit for Illumina. Libraries were quantified using the Agilent DNA 7500 Kit, and samples were mixed together at equimolar concentration. One biological replicate was sequenced using the Illumina HiSeq 2500 platform to generate 50 bp SE reads and carried out at the Tufts University Core Facility.

### PolyA-Selected RNA-Seq

Total RNA was harvested using the Ambion RNAqueous-4PCR kit and quantified using the Agilent RNA 6000 Nano kit. RNA-Seq libraries were generated from 2 µg RNA using NEBNext Poly(A) mRNA Magnetic Isolation Module and NEBNext Ultra Directional RNA Library Prep Kit for Illumina (NEB #E7490 and #E7420). Libraries were quantified using the Agilent DNA 7500 Kit, and samples were mixed together at equimolar concentration. Two biological replicates were sequenced using the Illumina HiSeq 2500 platform to generate 50 bp SE reads and carried out at the Tufts University Core Facility.

### Ribo-minus Total RNA-Seq

Total RNA was isolated from cells using TRIzol (Invitrogen), following manufacturer’s instructions. RNA was quantified using Agilent RNA 6000 Nano kit. ERCC spike-in controls (ThermoFisher) were added to 500 ng of total RNA and ribominus selection was performed using the NEBNext® rRNA Depletion Kit. RNA-Seq libraries were generated using the NEBNext Ultra Directional RNA Library Prep Kit for Illumina. Libraries were quantified using the Agilent DNA 7500 Kit, and samples were mixed together at equimolar concentration. One biological replicates was sequenced using the Illumina NextSeq550 platform to generate 75 bp PE reads and carried out at the Tufts University Core Facility.

### ChIP-Sequencing

ChIP-Seq was performed on mock-infected or infected MRC5 cells as described previously (11), with antibodies listed in Table 1. Libraries were quantified using the Agilent DNA 7500 Kit, and samples were mixed together at equimolar concentration. For each IP two biological replicates were sequenced, for input at least two biological replicates were sequenced using the Illumina HiSeq 2500 platform was used to generate 50 bp SE reads and carried out at the Tufts University Core Facility.

### ATAC-Sequencing

We adapted the protocol from Buenrostro et al. (2013), and previously published the data see SRA# PRJNA553559 (11).

### Data Availability

All datasets are publicly available at:

ATAC-Seq of WT (KOS) and ICP4 null (n12) HSV-1 Productive Infection in MRC5 cells SRA: PRJNA553559

ICP4 ChIP-Seq of WT (KOS) HSV-1 Productive Infection in MRC5 cells SRA: PRJNA553563

HSV-1 4SU-Seq SRA: PRJNA692715

HSV-1 POL3 Machinery-ChIP-Seq SRA: PRJNA693164 HSV-1 DM-tRNA-Seq SRA: PRJNA692681

HSV-1 GTF3C Complex-ChIP-Seq SRA: PRJNA732084

ChIP-Seq for POLR2A in HSV-1 infected human fibroblasts SRA: PRJNA732212

PolyA-Selected RNA-Seq in HSV-1 infected human fibroblast cells SRA: PRJNA732134 Total RNA-Seq in HSV-1 Infected Human Fibroblasts SRA: PRJNA732091

### Bioinformatic Analysis

Data was uploaded to the Galaxy web platform, and we used the public server at usegalaxy.org to analyze the data (76).

### Transcriptomic—DM-tRNA-Seq, 4SU-Seq, Ribo-minus Total RNA-Seq, PolyA-Selected RNA-Seq

We quantified mature and premature tRNA in 4SU-Seq and DM-tRNA-Seq datasets similar to the analysis described in (29). To discriminate between pre-and mature-tRNA species we first mapped sequencing data to an assembly of mature-tRNA sequences wherein 5’ and 3’ leader sequences are absent, introns are spliced, and 3’ CCA tails are present. Unaligned reads were then mapped to a modified host (hg38) genome in which tRNA genes were masked and instead appended as an additional chromosome containing pre-tRNA sequences with introns, and 5’ and 3’ leader sequences. Mature and premature-tRNA assemblies for hg38 were generated using the tRAX pipeline (77). Mitochondrial and nuclear tRNA loci were based on GtRNAdb (78, 79) and tRNAscan-SE (80) for hg38.

To assess mRNA levels, Ribominus Total or PolyA-selected RNA-Seq data was aligned sequentially using HISAT2 to the human genome (hg38) and HSV-1 strain KOS genome (KT899744.1) (81). FeatureCounts was performed using the KT89974.1 CDS as the reference GFF (82). For PolyA-selected RNA-Seq raw counts were normalized as mapped reads per billion total reads per kilobase (MR/BTR/Kb). For Ribominus Total RNA-Seq raw counts were normalized as mapped reads per million ERCC spike-in reads per kilobase (MR/MSI/Kb).

### Genomic—ChIP-Seq and ATAC-Seq

For ChIP-Seq, data was first aligned using Bowtie2 (83) to the human genome (hg38), and then unaligned reads were mapped to the HSV-1 strain KOS genome (KT899744.1). A modified version of the KT899744.1 genome was created, removing one copy of the repeat joint region (Δ1-9603, Δ125,845-126,977, Δ145,361-151,974). Bam files were visualized using DeepTools bamcoverage (84) with a bin size of 1 to generate bigwig files. Data was viewed in IGV viewer and exported as EPS files. Bigwig files were normalized for sequencing depth and genome quantity. Mapped reads were multiplied by the ‘norm factor’ which was calculated as the inverse of (Input cellular reads)/(Input cellular+viral mapped reads (TMR))×Billion sample TMR or (Input viral reads)/TMR× Million sample TMR. ChIP-Seq experiments were repeated for a total of 2-3 biological replicates. The normalized bigwig files were averaged between replicates. Heatmaps and gene profiles were generated using MultiBigwigSummary on normalized cellular bigwig files to all UCSC annotated mRNAs or high-confident tRNA loci from GtRNAdb (78, 79). Gene profiles and heatmaps were plotted using plotProfile and plotHeatmap (84).

For ATAC-Seq, data was first aligned using Bowtie2 (83) to the mitochondrial genome. Unaligned reads were sequentially mapped to the human genome (hg38), and the HSV-1 strain KOS genome (KT899744.1) with the following parameters: –no-unal –local – very-sensitive-local –nodiscordant –no-mixed –contain –overlap –dovetail –phred33. Bam files were visualized using DeepTools bamcoverage (84) with a bin size of 1 to generate bigwig files. Data was viewed in IGV viewer and exported as EPS files. Cellular bigwig files were normalized for sequencing depth (excluding mitochondrial mapped reads), the y-axes values are mapped reads per billion reads. The normalized bigwig files were averaged between two biological replicates. Heatmaps and gene profiles were generated using MultiBigwigSummary (84) on normalized cellular bigwig files to all UCSC annotated mRNAs. Gene profiles and heatmaps were plotted using plotProfile and plotHeatmap (231).

