## Supplemental Figures for "Manipulation of RNA Polymerase III by Herpes Simplex Virus-1"

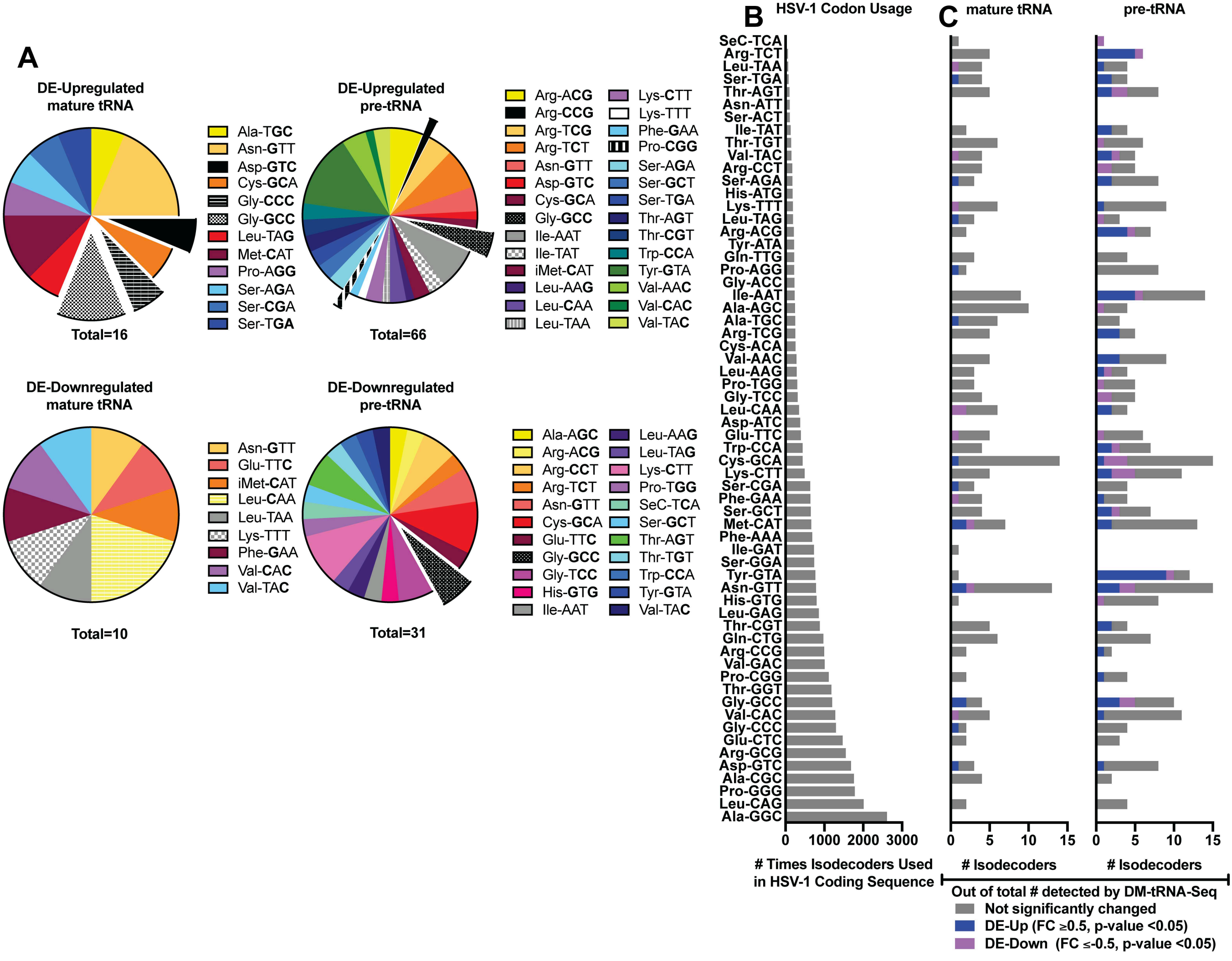

A
DE-Upregulated mature tRNA
Ala-TGC
Asn-GTT
Asp-GTC
Cys-GCA
Gly-CCC
Gly-GCC
Leu-TAG
Met-CAT
Pro-AGG
Ser-AGA
Ser-CGA
Ser-TGA
Total=16
DE-Upregulated pre-tRNA
Arg-ACG
Arg-CCG
Arg-TCG
Arg-TCT
Asn-GTT
Asp-GTC
Cys-GCA
Gly-GCC
Ile-AAT
Ile-TAT
iMet-CAT
Leu-AAG
Leu-CAA
Leu-TAA
Lys-CTT
Lys-TTT
Phe-GAA
Pro-CGG
Ser-AGA
Ser-GCT
Ser-TGA
Thr-AGT
Thr-CGT
Trp-CCA
Tyr-GTA
Val-AAC
Val-CAC
Val-TAC
Total=66
DE-Downregulated mature tRNA
Asn-GTT
Glu-TTC
iMet-CAT
Leu-CAA
Leu-TAA
Lys-TTT
Phe-GAA
Val-CAC
Val-TAC
Total=10
DE-Downregulated pre-tRNA
Ala-AGC
Arg-ACG
Arg-CCT
Arg-TCT
Asn-GTT
Cys-GCA
Glu-TTC
Gly-GCC
Gly-TCC
His-GTG
Ile-AAT
Leu-AAG
Leu-TAG
Lys-CTT
Pro-TGG
SeC-TCA
Ser-GCT
Thr-AGT
Thr-TGT
Trp-CCA
Tyr-GTA
Val-TAC
Total=31
B
HSV-1 Codon Usage
C
mature tRNA
pre-tRNA
SeC-TCA
Arg-TCT
Leu-TAA
Ser-TGA
Thr-AGT
Asn-ATT
Ser-ACT
Ile-TAT
Thr-TGT
Val-TAC
Arg-CCT
Ser-AGA
His-ATG
Lys-TTT
Leu-TAG
Arg-ACG
Tyr-ATA
Gln-TTG
Pro-AGG
Gly-ACC
Ile-AAT
Ala-AGC
Ala-TGC
Arg-TCG
Cys-ACA
Val-AAC
Leu-AAG
Pro-TGG
Gly-TCC
Leu-CAA
Asp-ATC
Glu-TTC
Trp-CCA
Cys-GCA
Lys-CTT
Ser-CGA
Phe-GAA
Ser-GCT
Met-CAT
Phe-AAA
Ile-GAT
Ser-GGA
Tyr-GTA
Asn-GTT
His-GTG
Leu-GAG
Thr-CGT
Gln-CTG
Arg-CCG
Val-GAC
Pro-CGG
Thr-GGT
Gly-GCC
Val-CAC
Gly-CCC
Glu-CTC
Arg-GCG
Asp-GTC
Ala-CGC
Pro-GGG
Leu-CAG
Ala-GGC
0 1000 2000 3000
### Times Isodecoders Used in HSV-1 Coding Sequence
0 5 10 15
### Isodecoders
0 5 10 15
### Isodecoders
Out of total # detected by DM-tRNA-Seq
Not significantly changed
DE-Up (FC ≥0.5, p-value <0.05)
DE-Down (FC ≤-0.5, p-value <0.05)

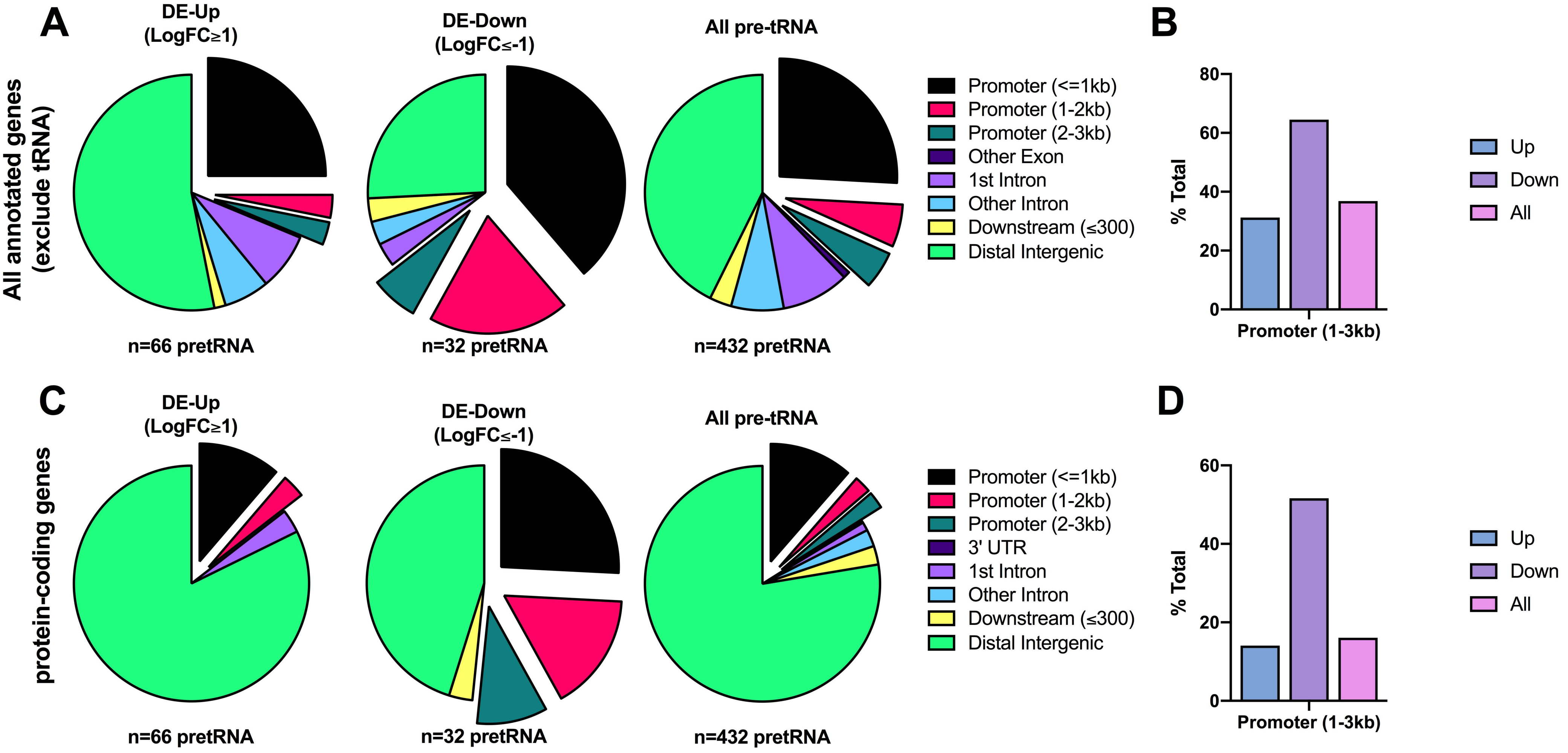

A
DE-Up (LogFC≥1)
DE-Down (LogFC≤-1)
All pre-tRNA
All annotated genes (exclude tRNA)
n=66 pretRNA
n=32 pretRNA
n=432 pretRNA
Promoter (<=1kb)
Promoter (1-2kb)
Promoter (2-3kb)
Other Exon
1st Intron
Other Intron
Downstream (≤300)
Distal Intergenic
B
% Total
0
20
40
60
80
Promoter (1-3kb)
Up
Down
All
C
DE-Up (LogFC≥1)
DE-Down (LogFC≤-1)
All pre-tRNA
protein-coding genes
n=66 pretRNA
n=32 pretRNA
n=432 pretRNA
Promoter (<=1kb)
Promoter (1-2kb)
Promoter (2-3kb)
3' UTR
1st Intron
Other Intron
Downstream (≤300)
Distal Intergenic
D
% Total
0
20
40
60
Promoter (1-3kb)
Up
Down
All

A

| Virus | Mutation | Mutation Type | Background Strain | tRNA upregulation |
|---|---|---|---|---|
| n12 | ΔICP4 | nonsense | KOS | Yes |
| d120 | ΔICP4 | deletion | KOS | Yes |
| d92 | ΔICP4/27 | deletion | KOS | Yes |
| 5dl1.2 | ΔICP27 | deletion | KOS | Yes |
| n212 | ΔICP0 | nonsense | KOS | Yes |
| d99 | ΔICP0 | deletion | KOS | Yes |
| n199 | ΔICP22 | nonsense | KOS | Yes |
| R3616 | ΔRL1 | deletion | F | Yes |
| F-ΔICP47 | ΔICP47 | deletion | F | Yes |

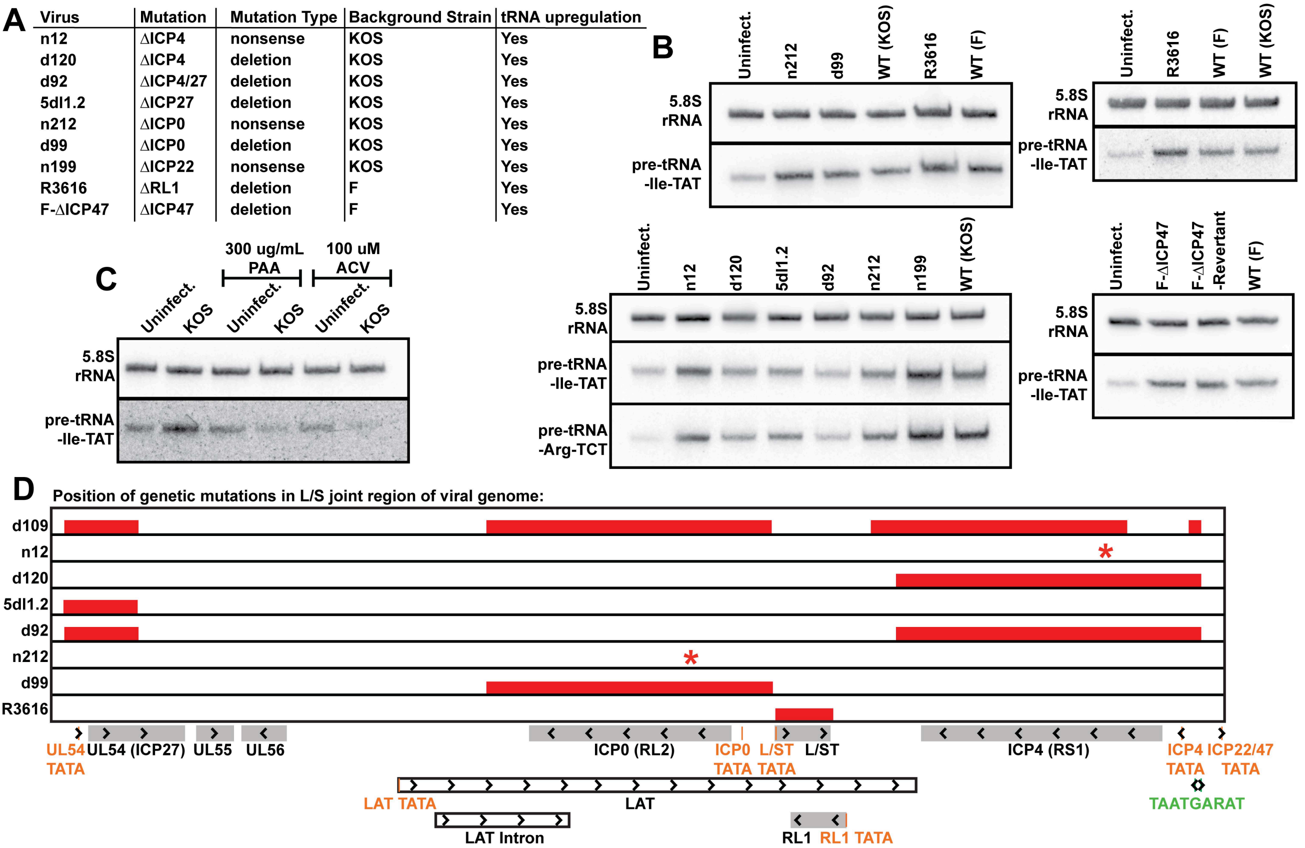

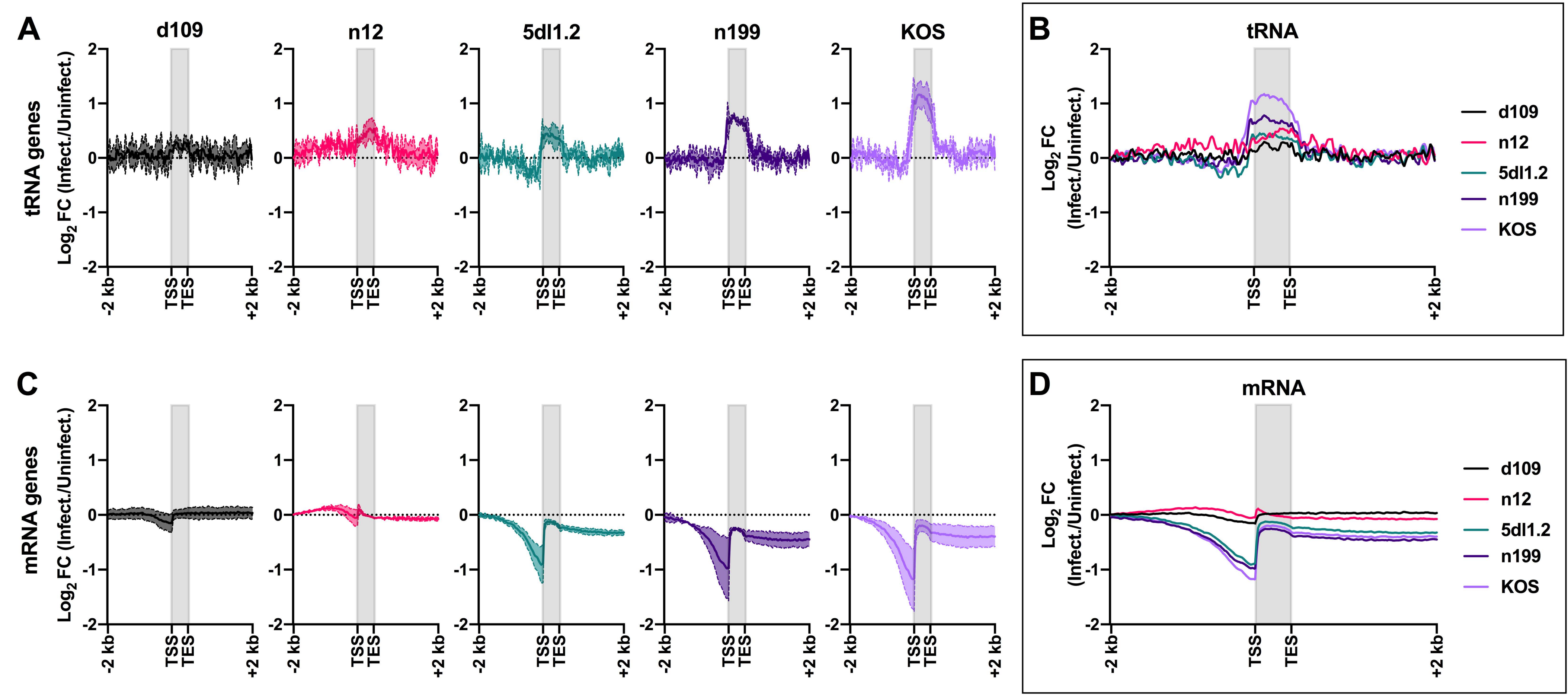

A
B
C
D
d109
n12
5dl1.2
n199
KOS
tRNA
mRNA
tRNA genes
mRNA genes
Log2 FC (Infect./Uninfect.)
-2 kb
TSS
TES
+2 kb
d109
n12
5dl1.2
n199
KOS

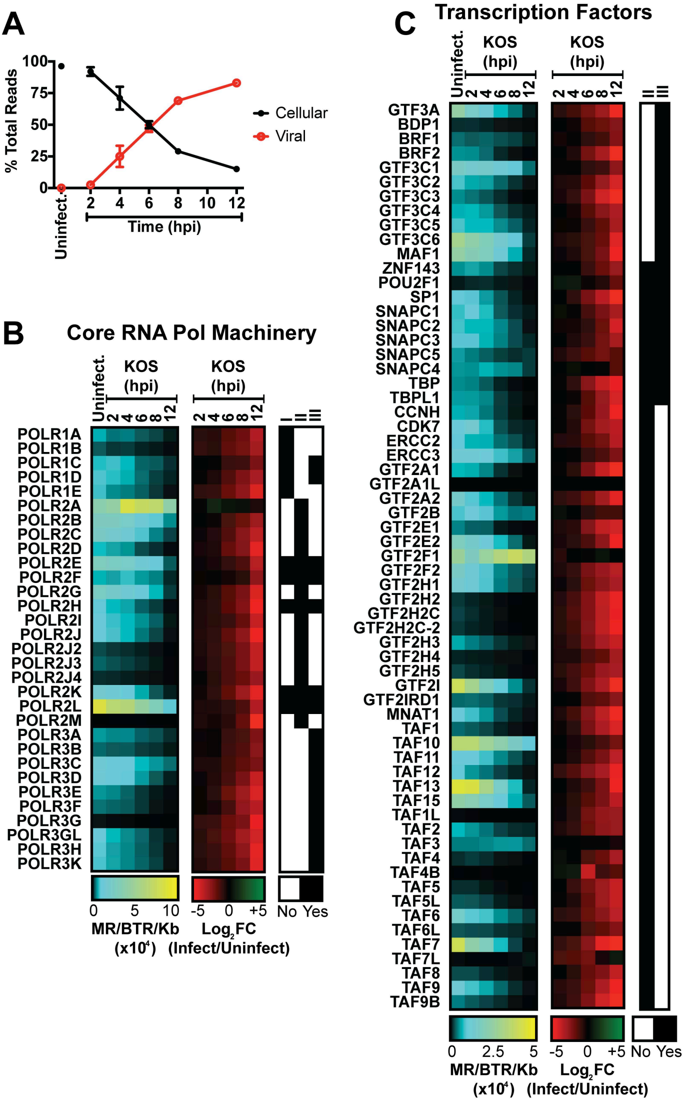

A
% Total Reads
100
75
50
25
0
Uninfect.
2
4
6
8
10
12
Time (hpi)
Cellular
Viral
B
Core RNA Pol Machinery
Uninfect.
KOS (hpi)
2 4 6 8 12
I II III
POLR1A
POLR1B
POLR1C
POLR1D
POLR1E
POLR2A
POLR2B
POLR2C
POLR2D
POLR2E
POLR2F
POLR2G
POLR2H
POLR2I
POLR2J
POLR2J2
POLR2J3
POLR2J4
POLR2K
POLR2L
POLR2M
POLR3A
POLR3B
POLR3C
POLR3D
POLR3E
POLR3F
POLR3G
POLR3GL
POLR3H
POLR3K
0 5 10
MR/BTR/Kb (x10^4)
-5 0 +5
Log2FC (Infect/Uninfect)
No Yes
C
Transcription Factors
GTF3A
BDP1
BRF1
BRF2
GTF3C1
GTF3C2
GTF3C3
GTF3C4
GTF3C5
GTF3C6
MAF1
ZNF143
POU2F1
SP1
SNAPC1
SNAPC2
SNAPC3
SNAPC5
SNAPC4
TBP
TBPL1
CCNH
CDK7
ERCC2
ERCC3
GTF2A1
GTF2A1L
GTF2A2
GTF2B
GTF2E1
GTF2E2
GTF2F1
GTF2F2
GTF2H1
GTF2H2
GTF2H2C
GTF2H2C-2
GTF2H3
GTF2H4
GTF2H5
GTF2I
GTF2IRD1
MNAT1
TAF1
TAF10
TAF11
TAF12
TAF13
TAF15
TAF1L
TAF2
TAF3
TAF4
TAF4B
TAF5
TAF5L
TAF6
TAF6L
TAF7
TAF7L
TAF8
TAF9
TAF9B
0 2.5 5
MR/BTR/Kb (x10^4)
-5 0 +5
Log2FC (Infect/Uninfect)
No Yes

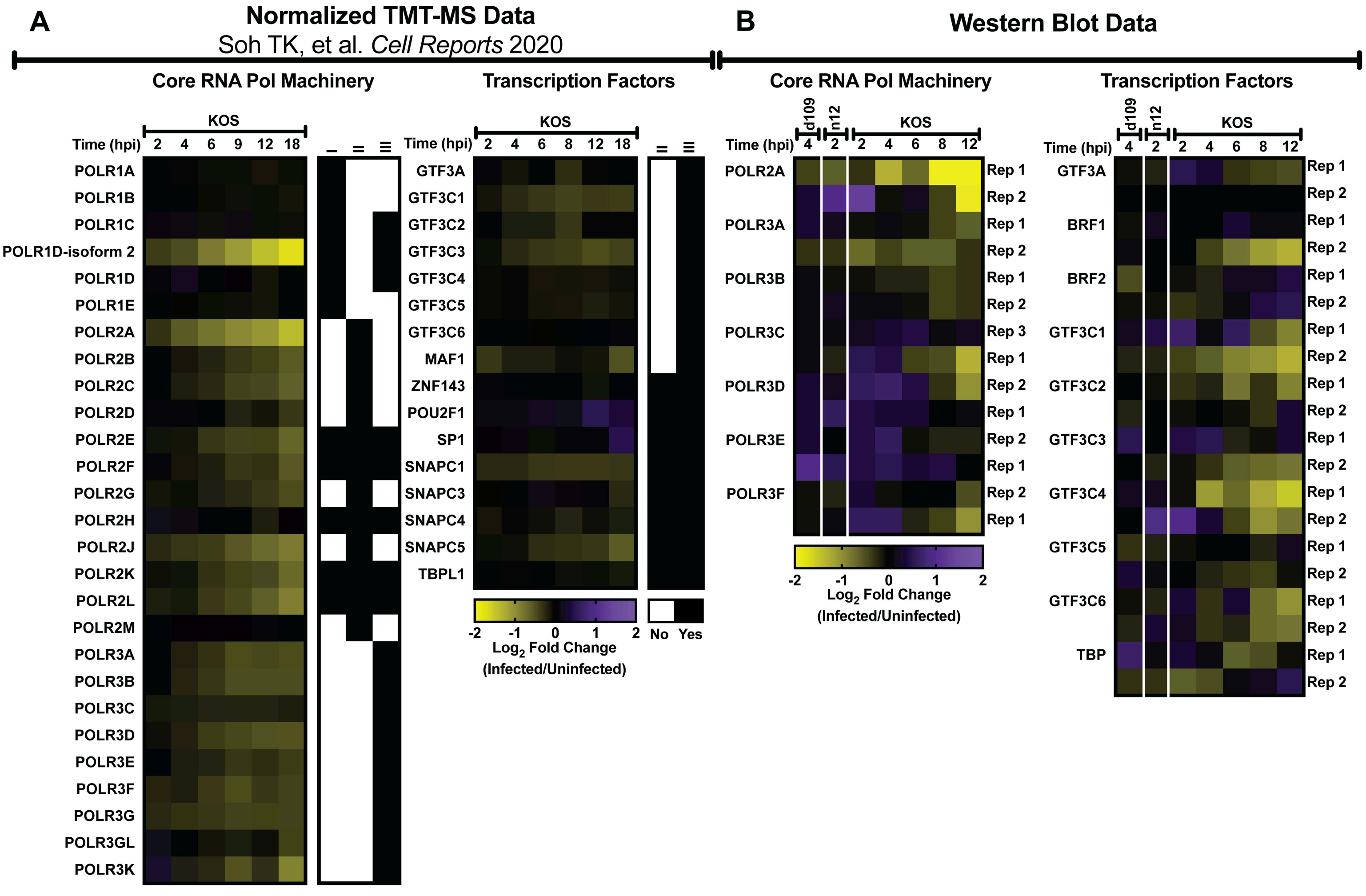
A
Normalized TMT-MS Data
Soh TK, et al. Cell Reports 2020
Core RNA Pol Machinery
Transcription Factors
KOS
Time (hpi) 2 4 6 9 12 18
POLR1A
POLR1B
POLR1C
POLR1D-isoform 2
POLR1D
POLR1E
POLR2A
POLR2B
POLR2C
POLR2D
POLR2E
POLR2F
POLR2G
POLR2H
POLR2J
POLR2K
POLR2L
POLR2M
POLR3A
POLR3B
POLR3C
POLR3D
POLR3E
POLR3F
POLR3G
POLR3GL
POLR3K
Time (hpi) 2 4 6 8 12 18
GTF3A
GTF3C1
GTF3C2
GTF3C3
GTF3C4
GTF3C5
GTF3C6
MAF1
ZNF143
POU2F1
SP1
SNAPC1
SNAPC3
SNAPC4
SNAPC5
TBPL1
-2 -1 0 1 2
Log2 Fold Change
(Infected/Uninfected)
No Yes
B
Western Blot Data
Core RNA Pol Machinery
Transcription Factors
d109 n12 KOS
Time (hpi) 4 2 2 4 6 8 12
POLR2A
POLR3A
POLR3B
POLR3C
POLR3D
POLR3E
POLR3F
Rep 1
Rep 2
Rep 3
-2 -1 0 1 2
Log2 Fold Change
(Infected/Uninfected)
GTF3A
BRF1
BRF2
GTF3C1
GTF3C2
GTF3C3
GTF3C4
GTF3C5
GTF3C6
TBP

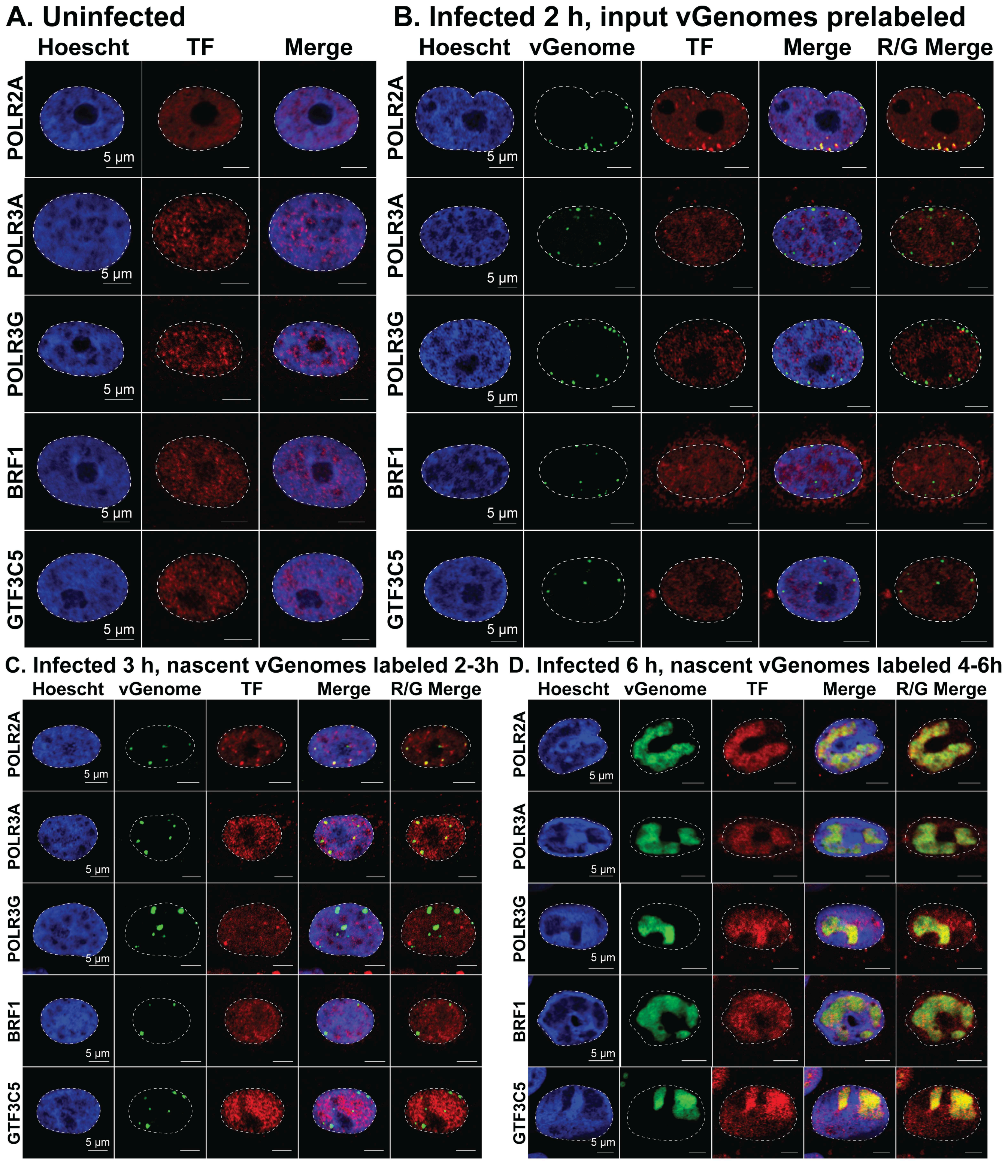
A. Uninfected
Hoescht
TF
Merge
B. Infected 2 h, input vGenomes prelabeled
Hoescht
vGenome
TF
Merge
R/G Merge
POLR2A
POLR3A
POLR3G
BRF1
GTF3C5
5 µm
C. Infected 3 h, nascent vGenomes labeled 2-3h
Hoescht
vGenome
TF
Merge
R/G Merge
D. Infected 6 h, nascent vGenomes labeled 4-6h
Hoescht
vGenome
TF
Merge
R/G Merge

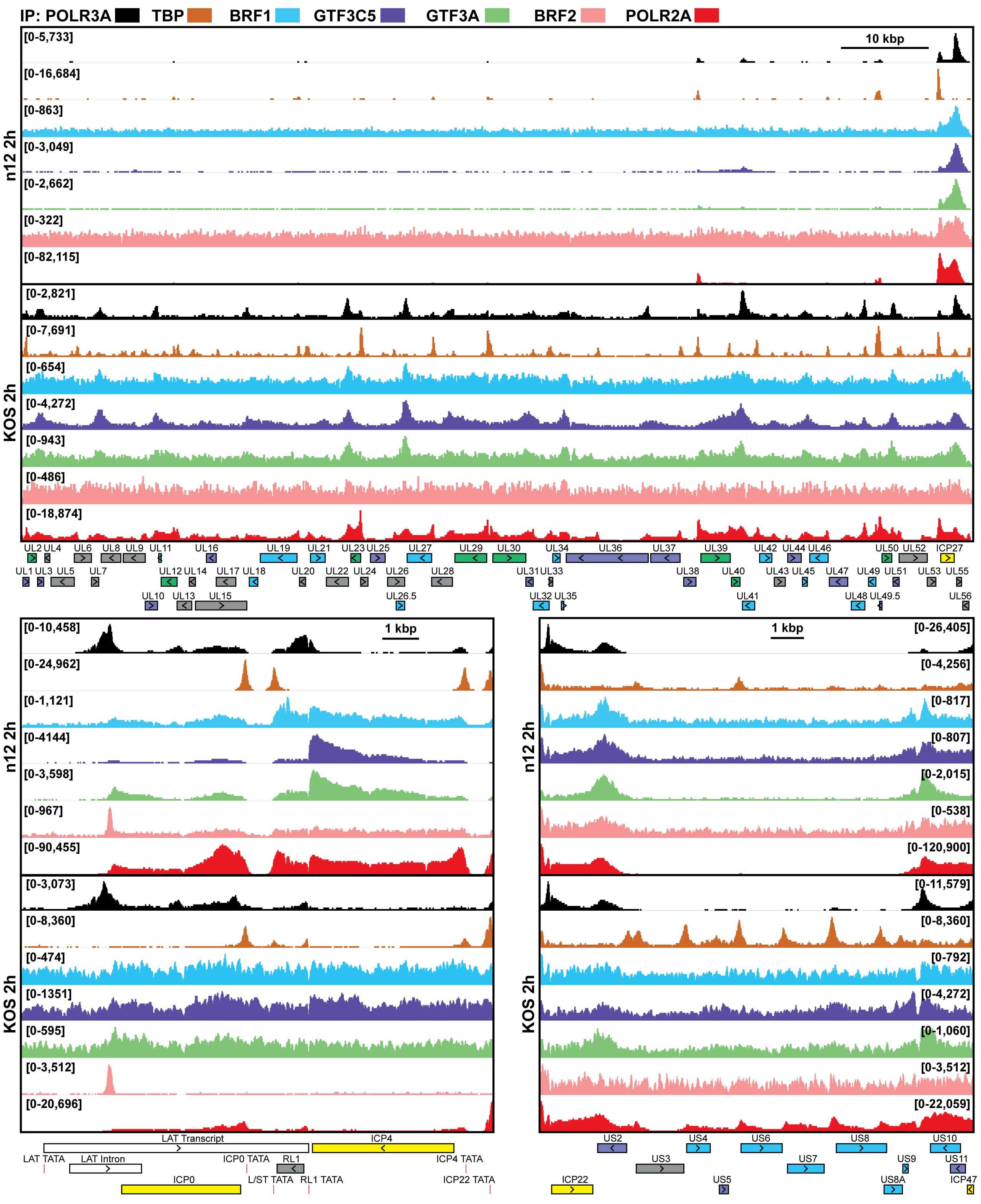
IP: POLR3A TBP BRF1 GTF3C5 GTF3A BRF2 POLR2A
n12 2h
KOS 2h
10 kbp
[0-5,733]
[0-16,684]
[0-863]
[0-3,049]
[0-2,662]
[0-322]
[0-82,115]
[0-2,821]
[0-7,691]
[0-654]
[0-4,272]
[0-943]
[0-486]
[0-18,874]
UL2 UL4 UL6 UL8 UL9 UL11 UL16 UL19 UL21 UL23 UL25 UL27 UL29 UL30 UL34 UL36 UL37 UL39 UL42 UL44 UL46 UL50 UL52 ICP27
UL1 UL3 UL5 UL7 UL12 UL14 UL17 UL18 UL20 UL22 UL24 UL26 UL28 UL31 UL33 UL38 UL40 UL43 UL45 UL47 UL49 UL51 UL53 UL55
UL10 UL13 UL15 UL26.5 UL32 UL35 UL41 UL48 UL49.5 UL56
1 kbp
[0-10,458]
[0-24,962]
[0-1,121]
[0-4144]
[0-3,598]
[0-967]
[0-90,455]
[0-3,073]
[0-8,360]
[0-474]
[0-1351]
[0-595]
[0-3,512]
[0-20,696]
LAT Transcript
ICP4
LAT TATA
LAT Intron
ICP0 TATA
RL1
ICP4 TATA
ICP0
L/ST TATA
RL1 TATA
ICP22 TATA
1 kbp
[0-26,405]
[0-4,256]
[0-817]
[0-807]
[0-2,015]
[0-538]
[0-120,900]
[0-11,579]
[0-8,360]
[0-792]
[0-4,272]
[0-1,060]
[0-3,512]
[0-22,059]
US2 US4 US6 US8 US10
US3 US7 US9 US11
ICP22 US5 US8A ICP47

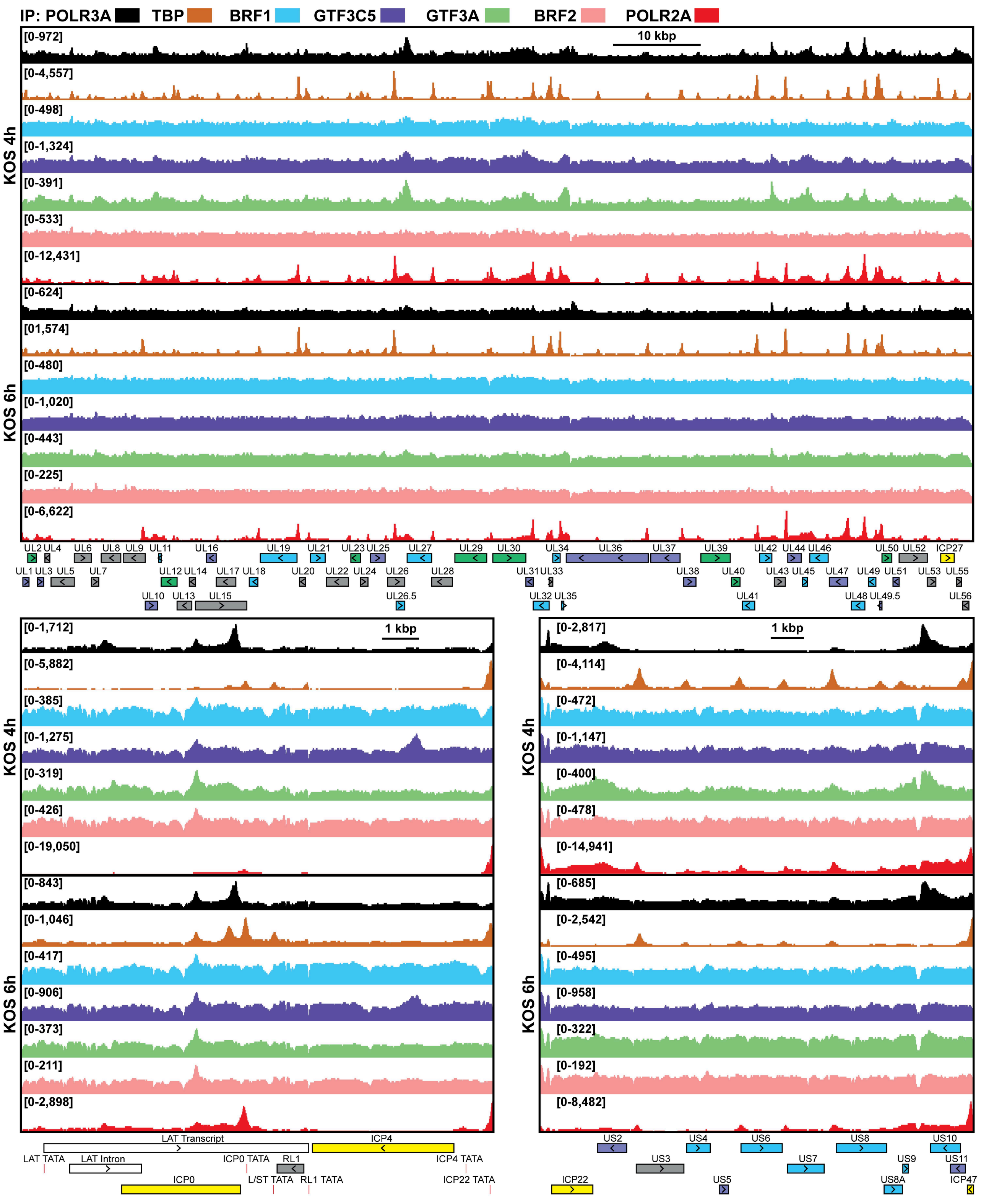
IP: POLR3A TBP BRF1 GTF3C5 GTF3A BRF2 POLR2A
10 kbp
KOS 4h
KOS 6h
[0-972]
[0-4,557]
[0-498]
[0-1,324]
[0-391]
[0-533]
[0-12,431]
[0-624]
[01,574]
[0-480]
[0-1,020]
[0-443]
[0-225]
[0-6,622]
UL2 UL4 UL6 UL8 UL9 UL11 UL16 UL19 UL21 UL23 UL25 UL27 UL29 UL30 UL34 UL36 UL37 UL39 UL42 UL44 UL46 UL50 UL52 ICP27
UL1 UL3 UL5 UL7 UL12 UL14 UL17 UL18 UL20 UL22 UL24 UL26 UL28 UL31 UL33 UL38 UL40 UL43 UL45 UL47 UL49 UL51 UL53 UL55
UL10 UL13 UL15 UL26.5 UL32 UL35 UL41 UL48 UL49.5 UL56
1 kbp
[0-1,712]
[0-5,882]
[0-385]
[0-1,275]
[0-319]
[0-426]
[0-19,050]
[0-843]
[0-1,046]
[0-417]
[0-906]
[0-373]
[0-211]
[0-2,898]
LAT Transcript
ICP4
LAT TATA
LAT Intron
ICP0 TATA
RL1
ICP4 TATA
ICP0
L/ST TATA
RL1 TATA
ICP22 TATA
[0-2,817]
[0-4,114]
[0-472]
[0-1,147]
[0-400]
[0-478]
[0-14,941]
[0-685]
[0-2,542]
[0-495]
[0-958]
[0-322]
[0-192]
[0-8,482]
US2 US4 US6 US8 US10
US3 US7 US9 US11
ICP22 US5 US8A ICP47

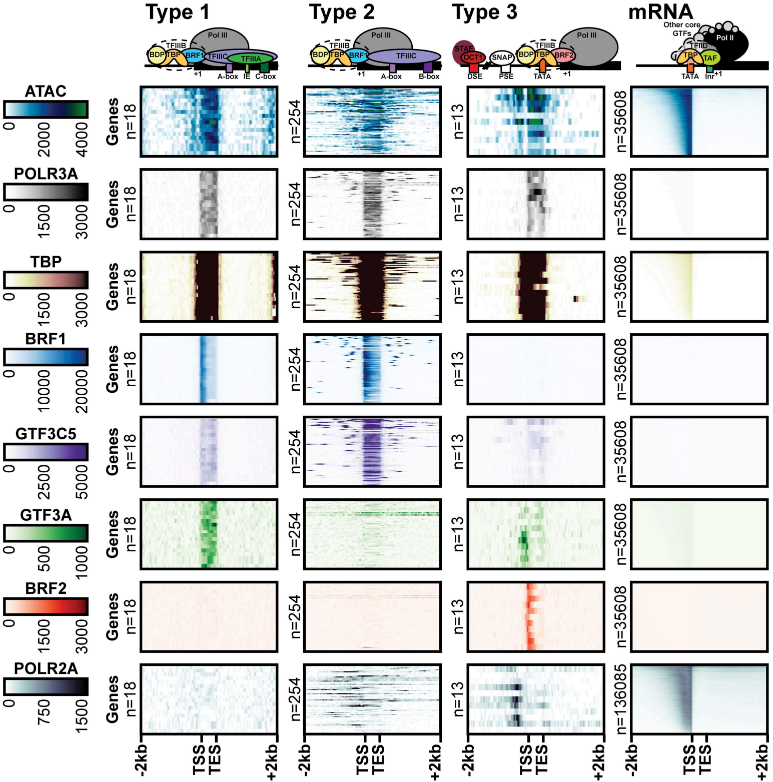

Type 1
Type 2
Type 3
mRNA
Pol III
TFIIIB
BDP
TBP
BRF1
TFIIIC
TFIIIA
A-box
IE
C-box
B-box
STAF
OCT1
SNAP
BRF2
DSE
PSE
TATA
+1
Other core GTFs
Pol II
TFIID
TAF
Inr
ATAC
0
2000
4000
POLR3A
1500
3000
TBP
BRF1
10000
20000
GTF3C5
2500
5000
GTF3A
500
1000
BRF2
POLR2A
750
Genes
n=18
n=254
n=13
n=35608
n=136085
-2kb
TSS
TES
+2kb

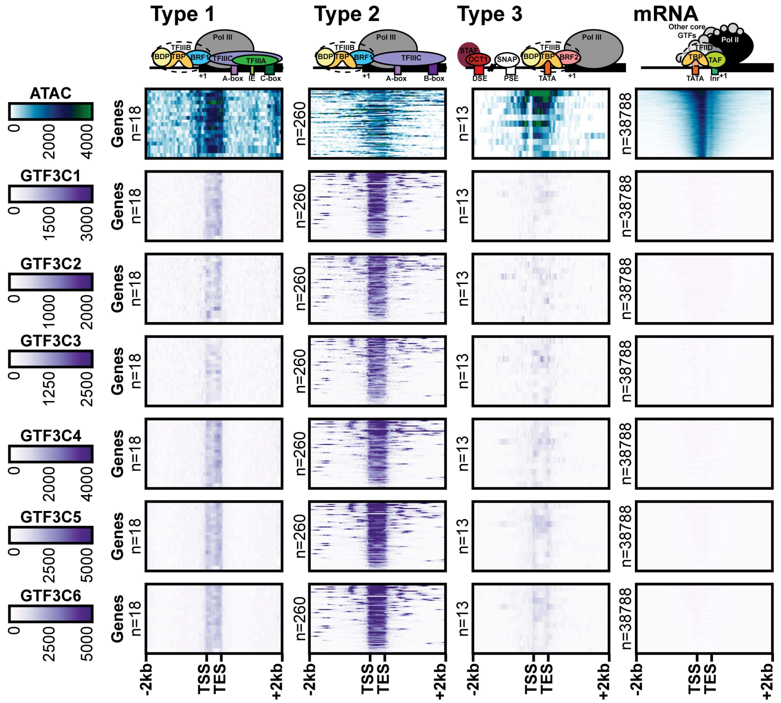

Type 1
Type 2
Type 3
mRNA
Pol III
TFIIIB
BDP
TBP
BRF1
TFIIIC
TFIIIA
+1
A-box
IE
C-box
B-box
STAF
OCT1
SNAP
BRF2
DSE
PSE
TATA
Other core GTFs
Pol II
TFIID
TAF
Inr
ATAC
0
2000
4000
GTF3C1
1500
3000
GTF3C2
1000
GTF3C3
1250
2500
GTF3C4
GTF3C5
5000
GTF3C6
Genes
n=18
n=260
n=13
n=38788
-2kb
TSS
TES
+2kb

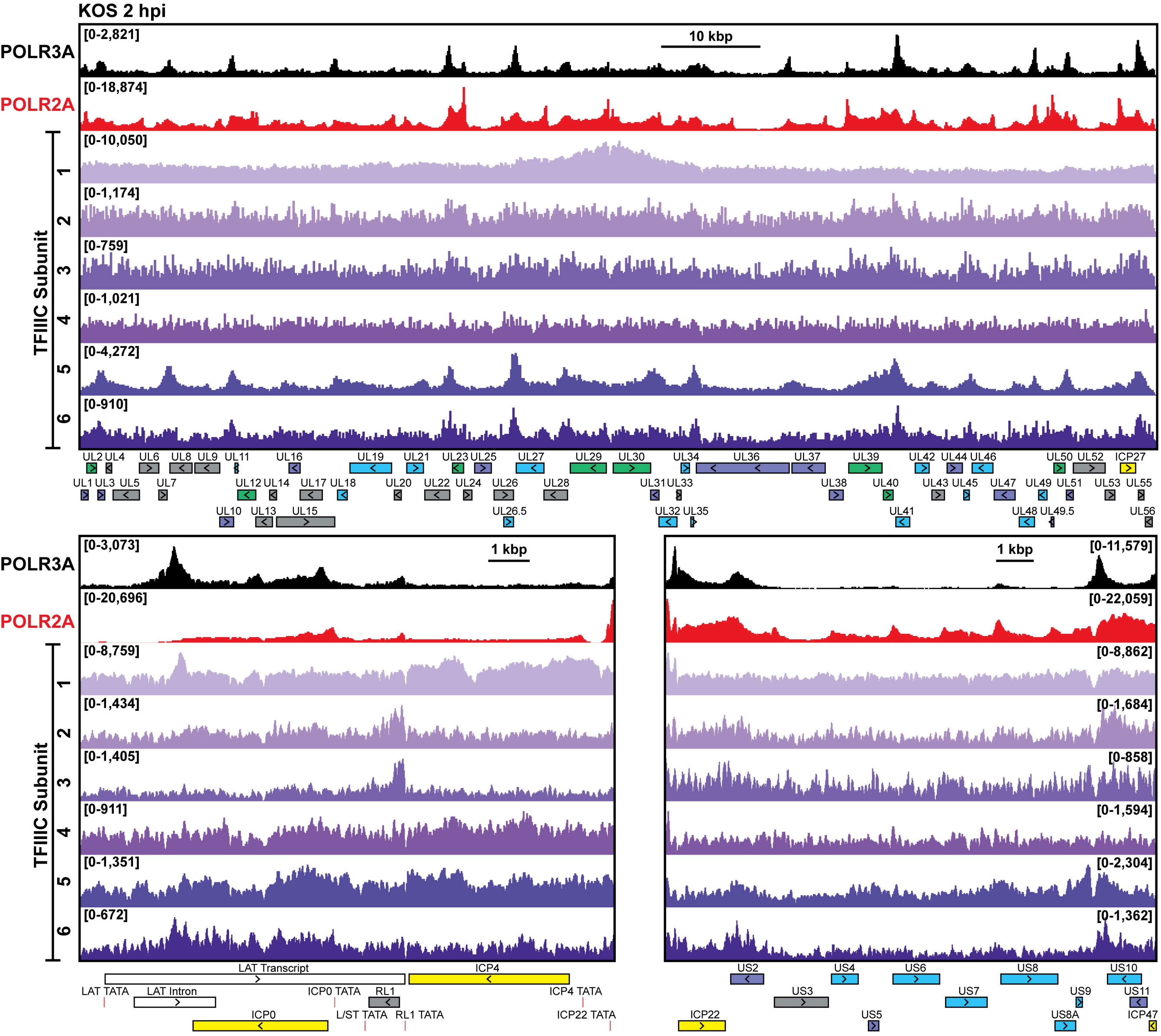

KOS 2 hpi
POLR3A
POLR2A
TFIIIC Subunit
1
2
3
4
5
6
10 kbp
[0-2,821]
[0-18,874]
[0-10,050]
[0-1,174]
[0-759]
[0-1,021]
[0-4,272]
[0-910]
UL2 UL4 UL6 UL8 UL9 UL11 UL16 UL19 UL21 UL23 UL25 UL27 UL29 UL30 UL34 UL36 UL37 UL39 UL42 UL44 UL46 UL50 UL52 ICP27
UL1 UL3 UL5 UL7 UL12 UL14 UL17 UL18 UL20 UL22 UL24 UL26 UL28 UL31 UL33 UL38 UL40 UL43 UL45 UL47 UL49 UL51 UL53 UL55
UL10 UL13 UL15 UL26.5 UL32 UL35 UL41 UL48 UL49.5 UL56
1 kbp
[0-3,073]
[0-20,696]
[0-8,759]
[0-1,434]
[0-1,405]
[0-911]
[0-1,351]
[0-672]
LAT Transcript
ICP4
LAT TATA
LAT Intron
ICP0 TATA
RL1
ICP4 TATA
ICP0
L/ST TATA
RL1 TATA
ICP22 TATA
[0-11,579]
[0-22,059]
[0-8,862]
[0-1,684]
[0-858]
[0-1,594]
[0-2,304]
[0-1,362]
US2 US4 US6 US8 US10
US3 US7 US9 US11
ICP22 US5 US8A ICP47
